# Targeting the Id1-Kif11-Aurka axis in triple negative breast cancer using combination therapy

**DOI:** 10.1101/760686

**Authors:** Reshma Murali, Binitha Anu Varghese, Nitheesh Karthikeyan, PT Archana, Wee Siang Teo, Andrea McFarland, Daniel L Roden, Holly Holliday, Christina Konrad, Aurelie Cazet, Eoin Dodson, Jason T George, Herbert Levine, Mohit Kumar Jolly, Alexander Swarbrick, Radhika Nair

**Author notes:** Equal first authors (Reshma Murali, Binitha Anu Varghese). Corresponding author: Dr Radhika Nair. Rajiv Gandhi Center for Biotechnology, Kerala, India.

## Abstract

Evidence points to breast cancer following a hierarchical model, with Cancer Stem Cells (CSCs) driving critical phenotypes of the bulk tumor. Chemoresistant CSCs are not an abstract concept but have clinical consequences as they drive relapse and ultimately lead to mortality in patients, making it imperative to understand how these subpopulations of cells survive. Our previous work (1-2) has demonstrated that the bHLH transcription factor, Inhibitor of Differentiation 1 (Id1) and it’s closely related family member Id3, have an important role in maintaining the CSC phenotype in the Triple Negative breast cancer (TNBC) subtype. A genetic screen conducted to further elucidate the molecular mechanism underlying the Id (Id1/3) mediated CSC phenotypes in TNBC revealed critical cell cycle genes such as Kif11 and Aurka as putative Id targets. We take this work forward by investigating how alteration in Kif11 and Aurka via Id proteins promotes the CSC phenotype in TNBC. Cells lacking Id are poised in a state of G0/G1 arrest from which they can re-enter the cell cycle. Intriguingly, depletion of Kif11 and Aurka independently did not phenocopy the G0/G1 arrest observed in Id knockdown (Id KD) cells. We have further explored the hypothesis that we can deplete the chemo resistant Id expressing CSC population by combining chemotherapy with targeted therapy using existing small molecule inhibitors (against Id target Kif11) to more effectively debulk the entire tumor. This work opens up exciting new possibilities of targeting Id targets like Kif11, in the TNBC subtype which is currently refractory to chemotherapy.

## Introduction

Breast cancer is a heterogeneous disease with different molecular subtypes displaying distinct pathological-clinical outcomes that have been successfully exploited in the management of the disease (3). The TNBC subtype does not express molecular markers such as ER and Her2 that are the basis of targeted therapies in other molecular subtypes of breast cancer (4). Consequently patients presenting with TNBC are left with few therapeutic choices, resulting in lower five-year survival rates when compared to the other subtypes (4-5). There is hence an urgent need to understand the molecular basis of TNBC in order to identify new drug targets.

The critical role of a subpopulation of cells termed Cancer Stem Cells (CSCs) in self-renewal, chemoresistance and metastasis has assumed great clinical importance in breast cancer (6-7). The Inhibitor of differentiation (Id) proteins are negative regulators of the basic helix-loop-helix (bHLH) transcription factors. The Id proteins are important for maintaining the CSC population and therefore tumour progression in TNBC(8) We have previously shown that Id1/3 (collectively known as Id) are critical for the CSC associated phenotypes in the TNBC molecular subtype(1). A detailed genetic screen analysis of Id knock down (Id KD) and Id1 expression models led to the identification of Kif11 and Aurka as putative Id targets.

The detailed mechanism by which Id controls the cell cycle is not clear, although Id is known to impact the pathway via decreased expression of Cyclins D1 and E, reduced phosphorylation of Rb as well as reduced Cyclin E-Cdk2 activity (9). In this work, we show how Id acts as a central focal point to coordinate the cell cycle genes Kif11 and Aurka and demonstrate that Id KD leads to cell cycle arrest in the G0/G1 phase of the cell cycle. Interestingly, we found that the depletion of Kif11 and Aurka independently did not phenocopy the G0/G1 arrest we observed in Id KD cells. We demonstrate that Id KD puts the brakes on the cell cycle resulting in a state of arrest at the G0/G1 phase via impacting cell cycle molecules and Id is a critical driver of self-renewal acting via Kif11 and Aurka. We found that the Id expressing tumor cells are resistant to chemotherapy, which forms the first line of treatment in TNBC. Interestingly, treatment with the Ispinesib, a small molecule inhibitor against Kif11 resulted in the reduced expression of Id in these cells. We finally exploit this finding to treat tumor cells with the chemotherapy Paclitaxel combined with Ispinesib to ablate the Id expressing chemo resistant tumor cells along with bulk tumor cells leading to more effective therapeutic targeting in the TNBC subtype.

## Results

### Id depletion leads to a G0/G1 cell cycle arrest which is reversible

It has been previously demonstrated that Id KD significantly affects pathways associated with cell cycle progression ((10-11). We first sought to validate this observation in our model system using the pSLIK (single lentivector for inducible knockdown) construct(12-13). We used the metastatic 4T1 cell line as it is representative of the TNBC subtype and Id proteins have been shown to play an important role in tumorigenesis in TNBC subtype. As reported before, we observed a significant decrease in the proliferative capacity of cells upon Doxycycline (Dox) induced Id KD in comparison to control conditions.

As proliferation is inextricably linked to the cell cycle, we next characterized the effect of Id KD on cell cycle progression. We found that Id depletion prevented cells from entering the S phase with accumulation in the G0/G1 phase, as seen in a significant increase in the G0/G1 fraction when compared to the controls (Figure 1A, B, C). To further elucidate the molecular mechanism through which Id controls the cell cycle, we analysed the effect of Id KD on the expression of key cell cycle genes which are vital at different phases of the cell cycle. The down regulation of Id significantly decreased the expression of Ccna2, Ccnb1, Ccnb2, Cdk1 and c-Myc as shown in Figure 1D. Interestingly, it shows an inverse correlation with Rb and p21, which are the negative regulators of these cell cycle genes (Figure1E).

**Figure 1.**
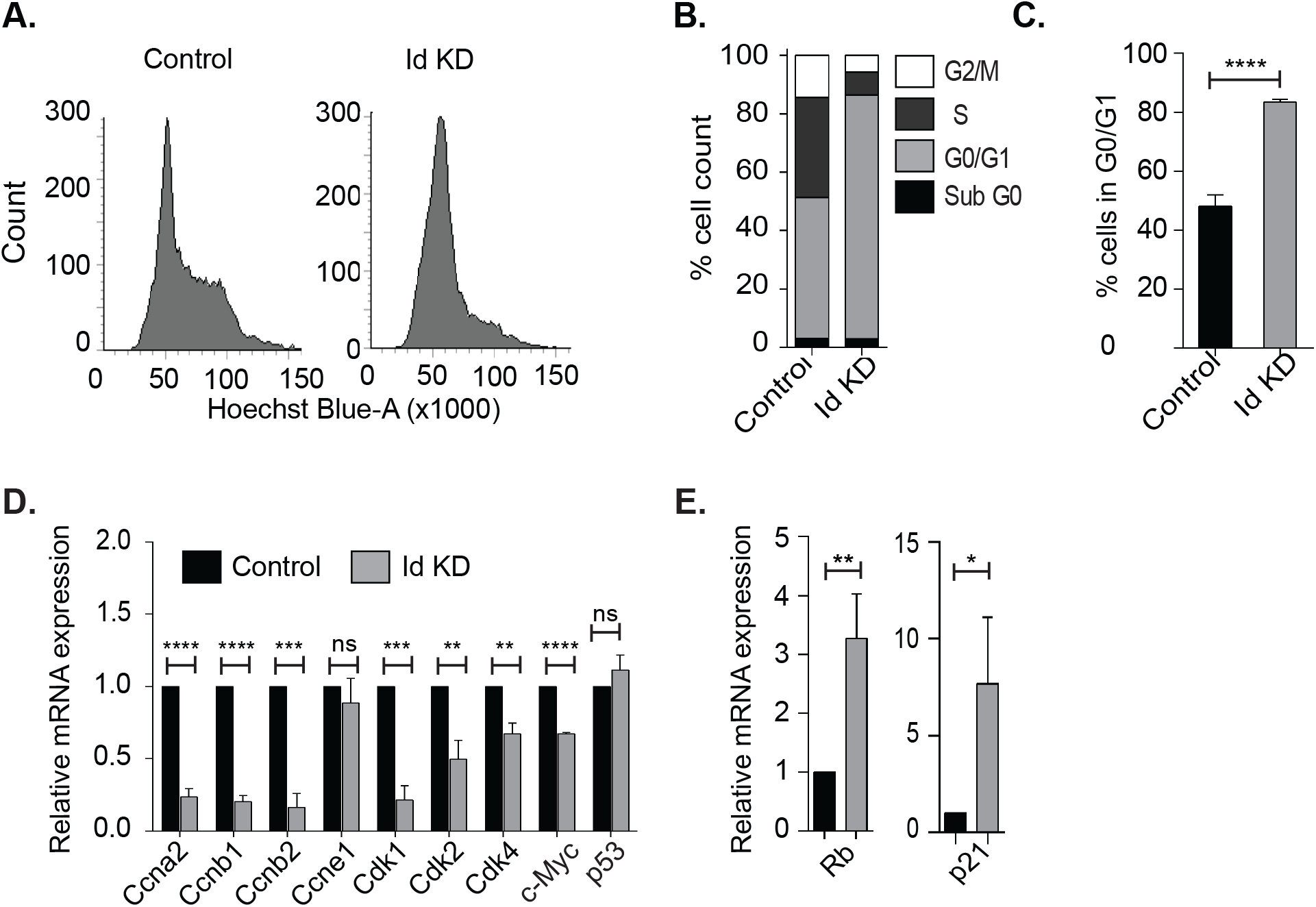
Effect of Id knockdown on cell cycle genes. (A) Flow cytometric analysis of cell cycle on Control and Id KD cells after labelling with Hoechst 33342 stain. (B) Comparing the percentage cell count in each phase of the cell cycle showing a significant increase in the G0/G1 phase in Id KD conditions. (C) Percentage cell count in G0/G1 phase of Control and Id KD cells. (D)The relative mRNA expression level of cell cycle genes Ccna2, Ccnb1,Ccnb2, Ccne1, Cdk1, Cdk2, Cdk4, c-Myc and p53 in Control and Id KD cells quantified using qRT-PCR. (E) Relative mRNA expression of Rb and p21 in Id KD cells with respect to Control cells were quantified using qRT-PCR. Data were normalized to beta-actin and analyzed by the 2−ΔΔCt method.

We have already demonstrated that Id KD significantly compromises other key CSC phenotypes like self renewal and migration (10). As Id marks a CSC in the TNBC subtype, we next looked at the expression of the CSC markers CD24 and CD29 in the Id KD system (10, 14-17). Contrary to our hypothesis, we found that the percentage of cells marked by CD29+/CD24+ in the Id KD population is significantly higher in comparison to the controls (Supplementary Figure1A). This suggests that CD29+/CD24+ does not mark the CSCs in 4T1 model system unlike other model systems (18). Therefore Id depletion clearly affects the key CSC phenotypes such as proliferation, self-renewal and migration which are closely linked to the cell cycle in the TNBC subtype.

### Identification of putative Id regulated genes impacting on the cell cycle

To characterize the network of genes regulated by Id proteins, functional annotation analysis was performed on gene array and RNA sequencing data from two different TNBC models of tumour cells marked by either Id depletion or Id1 expression, as described previously (1) (Figure 2A). The Id1 expression model analysed genes whose expression was associated with Id1, whereas the Id depletion model attempted to identify downstream targets of Id proteins.

**Figure 2.**
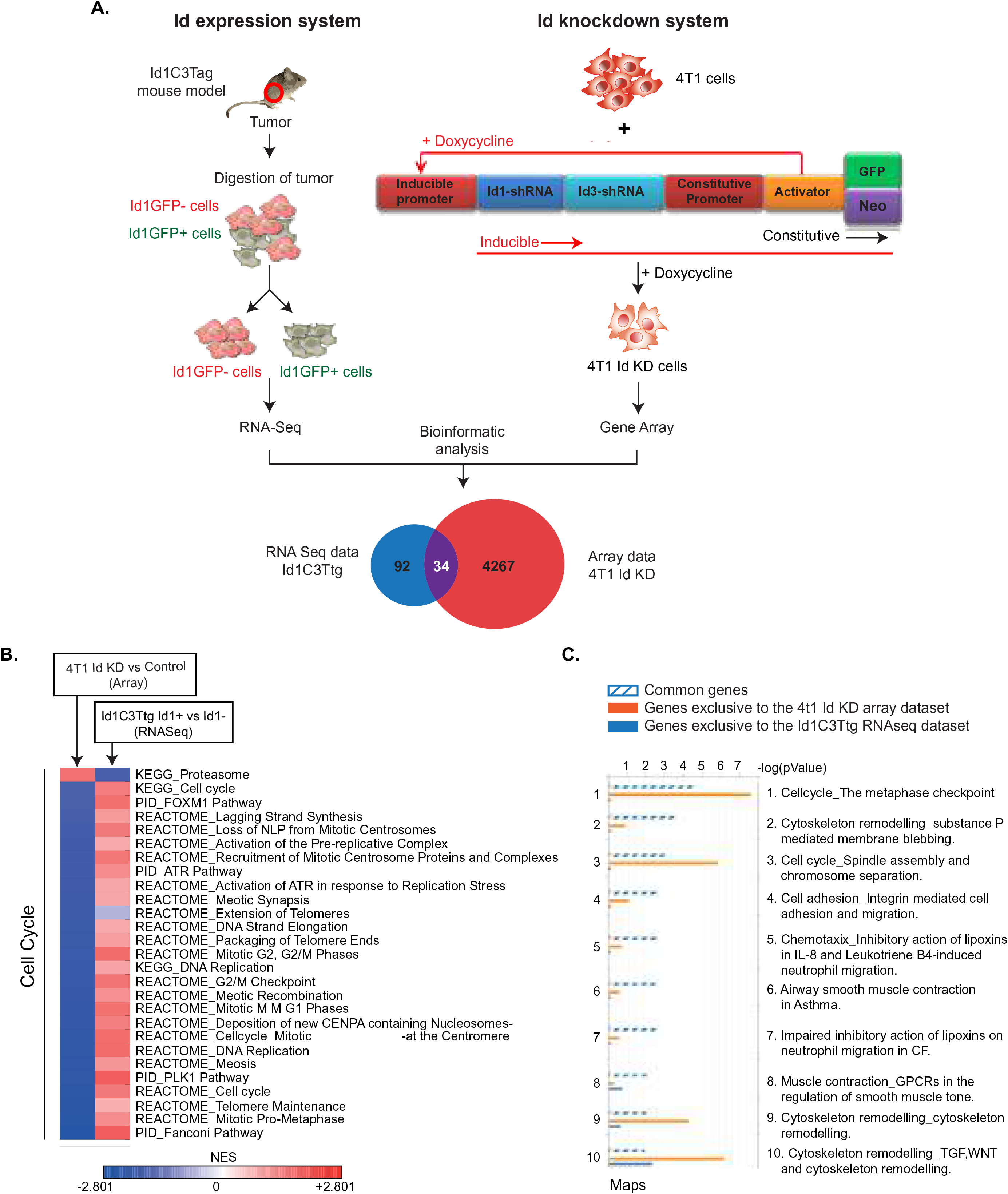
Identification of putative ID regulated genes. (A)To characterize the network of genes regulated by Id proteins, functional annotation analysis was performed on gene array and RNA sequencing data from two different TNBC models of tumour cells marked by either Id depletion or expression. Aiming to discover high confidence genes involved in the Id gene regulatory network, lists of differentially expressed genes between the Id expressing or depleted TNBC models and their controls were compared using MetaCoreTM software. By comparing these two datasets, lists of MetaCore network objects common to both experiments as well as those unique to each of the two data sets was generated The data was uploaded in MetaCore and filtered using an adjusted p-value threshold of 0.05, resulted in 4301 network objects that were differentially expressed between the Id KD and the 4T1 control cells. Similarly the Id1C3Ttg RNA Sequencing data revealed 126 network objects specific to mouse differentially expressed between Id+ and Id-cells. When comparing these lists of network object for the two TNBC models, 34 high confidence MetaCore network objects were identified as common to both data sets of differently expressed genes. Finally, the genes were mapped to network objects in MetaCore, resulting in a list of 34 genes that were significantly regulated in both models of TNBC. (B) To characterize the network of genes regulated by Id, enrichment analysis was performed on the 34 candidate genes and visualized by process networks and pathway maps. The enriched pathways were involving cytoskeleton remodelling, integrin-mediated cell adhesion and migration, and chemotaxis, which are all key steps in EMT and metastasis. (C) Id expression showed enrichment for pathways involving hypoxia-induced epithelial-mesenchymal transition, WNT pathway in development, cytoskeleton remodelling and cell cycle.

To study the phenotypes associated with depletion of Id as well as to assess its downstream targets, the gene expression profile of three independent replicates of control and Id KD cells was compared by microarray analysis to generate a list of differentially expressed genes between Id depleted and control cells (Supplementary Table 1). In addition, we used the Id1C3Tag model system to prospectively isolate Id1+ cells as described earlier(1). The gene expression profiles of the Id1+ and Id1-cells from three independent Id1C3Tag tumours were compared by RNA sequencing. This resulted in a list of differentially expressed genes between the Id+ and Id-mouse TNBC cells (Supplementary Table 2). To characterize the network of genes regulated by Id, enrichment analysis was performed on the candidate genes and visualized by process networks and pathway maps. Interestingly, when we looked at the pathway analysis generated from the differentially expressed gene lists of both models, cell cycle pathways were among the top hits (Supplementary Figure 1B, C)

Aiming to discover high confidence genes involved in the Id gene regulatory network, lists of differentially expressed genes in the Id depleted TNBC model and their controls were compared using MetaCoreTM software. The data was uploaded in MetaCore and filtered using an adjusted p-value threshold of 0.05, resulted in 4301 network objects that were differentially expressed between the Id KD and the control cells (Figure 2A). Similarly the Id1C3Tag RNA Sequencing data revealed 126 network objects differentially expressed between Id1+ and Id1-cells (Figure 2A). By comparing these two datasets, lists of MetaCore network objects common to both experiments as well as those unique to each of the two data sets were generated (Supplementary Figure 1D).When comparing these lists of network objects from the two TNBC models, 34 high confidence MetaCore network objects were identified as common to both the data sets of differently expressed genes. Finally, the genes were mapped to network objects in MetaCore, resulting in a list of 26 genes that were significantly regulated in both models of TNBC (Table 1).

**Table 1.**
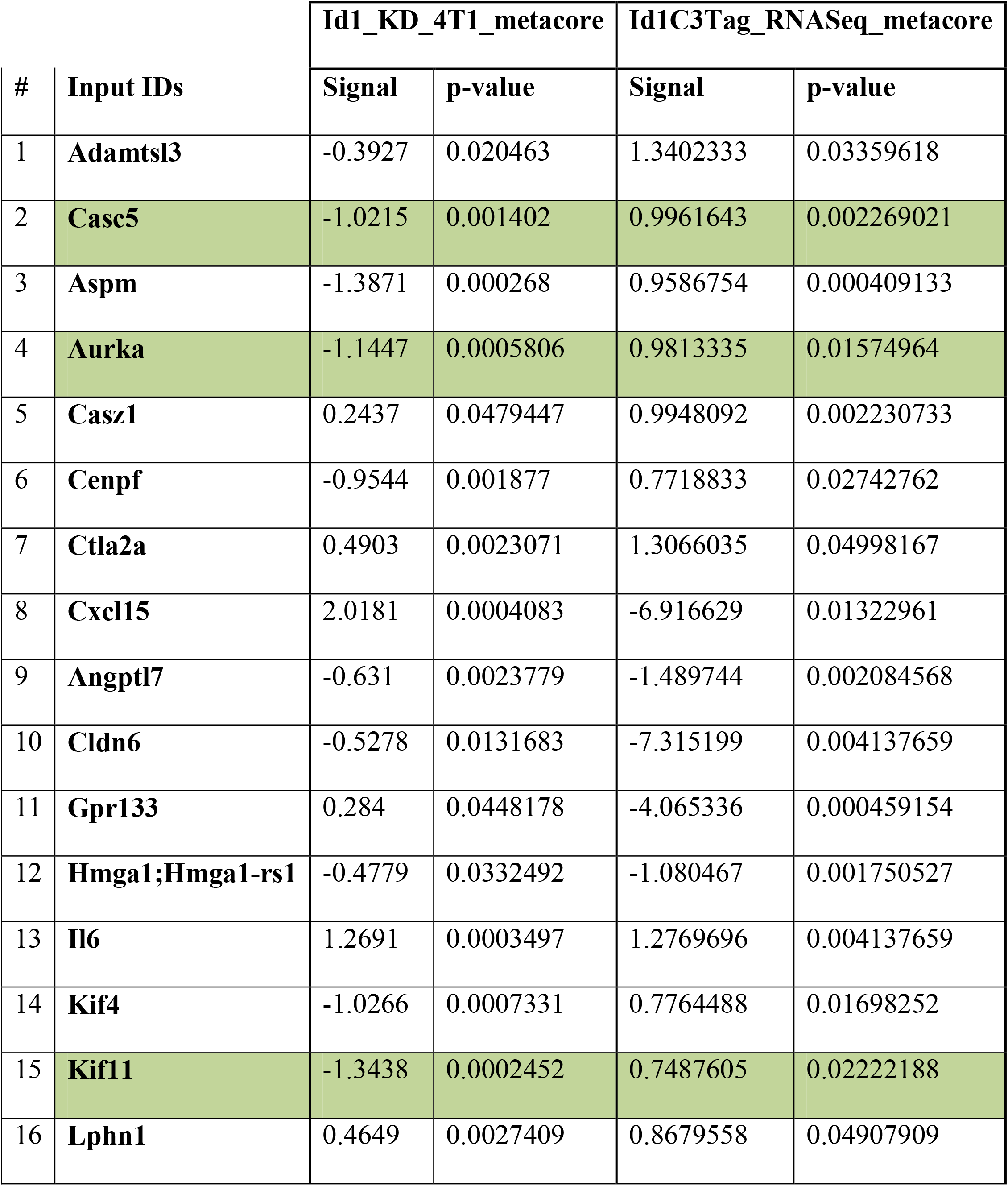
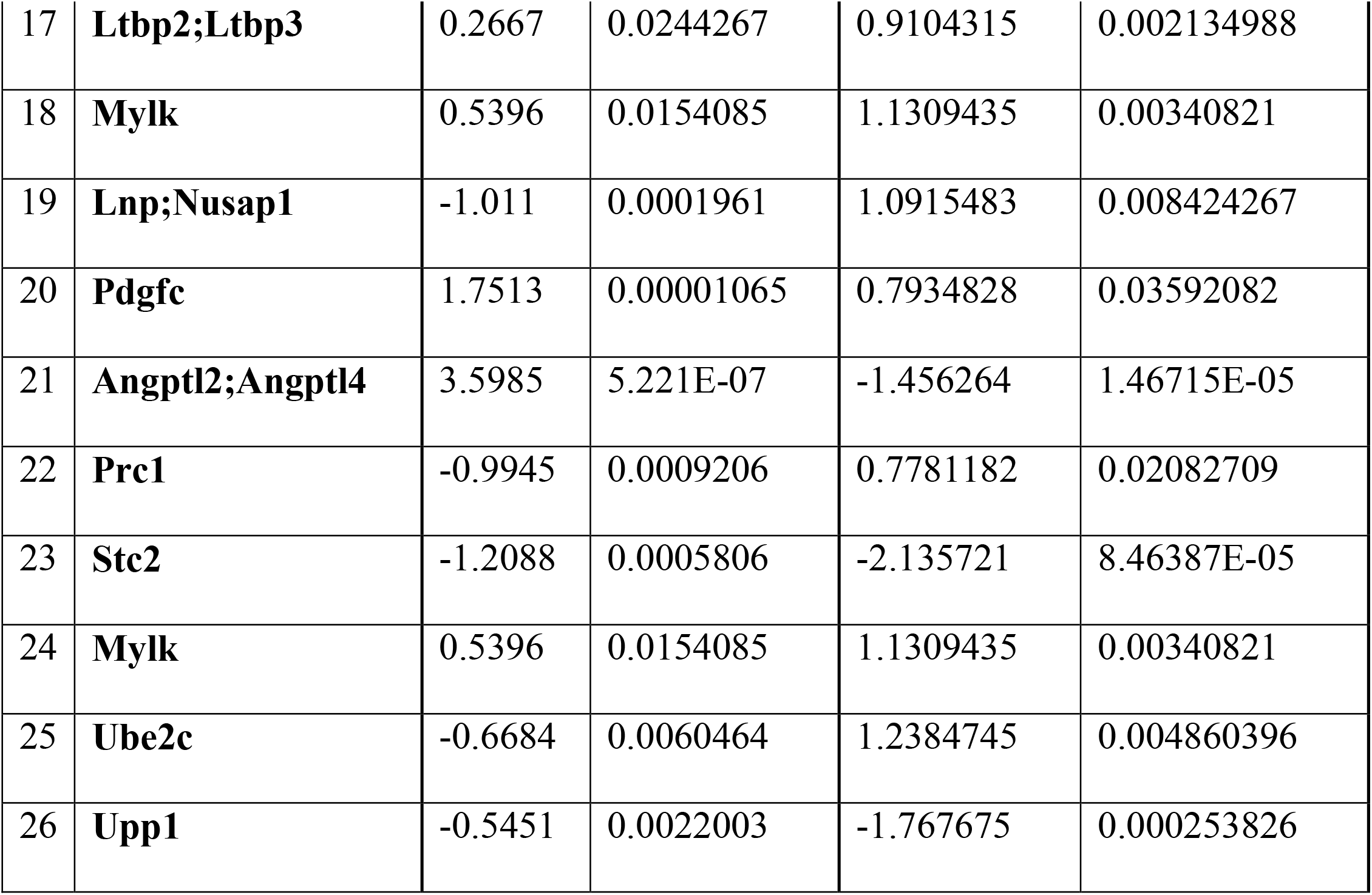
List of differentially expressed genes common to both the models.

We first analysed the pathways controlled by the 26 putative Id targets. Interestingly, the main pathways regulated by Id involved the cell cycle, specifically the metaphase checkpoint, spindle assembly and chromosome segregation (Figure 2B, C). Strict regulation of cell cycle progression and proliferation is essential for normal cellular function. Disruption of checkpoint control and aberrant regulation of the cell cycle are thus observed in tumourigenesis resulting in uncontrolled cell proliferation. A key function of Id is the stimulation of cell cycle progression and proliferation by controlling the activity of cell cycle regulators. Studies already reported that over expression of Id has been associated with up regulated cell cycle progression in tumourigenesis (9, 11, 19-20). Also enriched were pathways involving cytoskeleton remodelling, integrin-mediated cell adhesion and migration and chemotaxis, which are all key steps in epithelial to mesenchymal transition (EMT) and metastasis (Figure 2C). Analysis of each individual experiment, along with the genes common to both data sets, showed a similar result with Id depletion affecting mainly the cell cycle pathway, DNA damage, checkpoints, and cytoskeleton remodelling. Id expression model alone showed enrichment for pathways involving hypoxia-induced epithelial-mesenchymal transition, WNT pathway in development, cytoskeleton remodelling and cell cycle (Figure 2C).

We next investigated the process networks that were significantly enriched by the genes common to both the Id1 expressing and Id depleted models. This analysis identified process networks involved in cell cycle, cell adhesion and the cytoskeleton but also regulation of angiogenesis and inflammation. Casc5, Aspm, Aurka, Cenpf, Dynamin, Kif11, Kif4a, and Ube2c were the main drivers of the cell cycle and cytoskeleton networks, whereas the process networks involved in regulation of angiogenesis were enriched for Ctgf, Il6, Angptl4, and Oxtr. Process networks involved in cell adhesion were enriched for Il6, Pdgfc, and Ctgf. In addition, DNA damage checkpoint and mismatch repair (MMR) as well as regulation of EMT were also enriched in the Id depleted model whereas Id1 expression uniquely affected cell-matrix adhesion and interactions (Figure 2C).

We looked at the EMT program which is an important driver of the CSC state and found a significant change in the E cadherin and Vimentin protein levels (Supplementary Figure 2A, B). We next looked at the EMT state using a bioinformatics approach and interestingly found that the EMT scores of these samples using an inferential scoring metric(21) did not show any significant change in the EMT scores, indicating that the EMT status of the cells was not altered upon Id KD (Supplementary Figure 2C). This clearly suggests as elaborated in the discussion section that the CSC phenotype was being driven by Id using a unique slew of genes.

### Identification of Kif11 and Aurka as potential Id targets in disrupting the cell cycle pathway

Among the 26 differentially expressed genes common to the two TNBC models marked by Id depletion or Id1 expression, 16 genes were prioritized for validation. These were chosen based on a significant p value (< 0.05) and at least 1.5 fold (0.58 log2 FC) down or up regulation compared to the controls. Most of these genes had opposite regulation in the two TNBC models which was consistent with the fact that one model was marked by Id depletion whereas the other was an Id1 expression model. In addition, 8 potential cancer stem cell markers and genes previously implicated in Id biology were chosen based on cell surface localization, significant enrichment in Id+ cells and availability of antibodies. Altogether 60 candidate genes were identified for further validation as putative Id candidate target genes (Supplementary Table 3).

As Id functions as a key regulator of cell cycle progression and unlimited proliferation is a key phenotype of cancer cells, the effect of target gene knockdown on the proliferative phenotype of the Id KD cells was assessed for the 61 candidate genes identified using a targeted siRNA screen. Interestingly, the target genes that showed the greatest effect on the viability and thus the proliferative phenotype of the Id KD cells with more than 50% were Kif11, Casc5, Ccnd1 and Aurka (Figure 3A, Supplementary Table 3).

**Figure 3.**
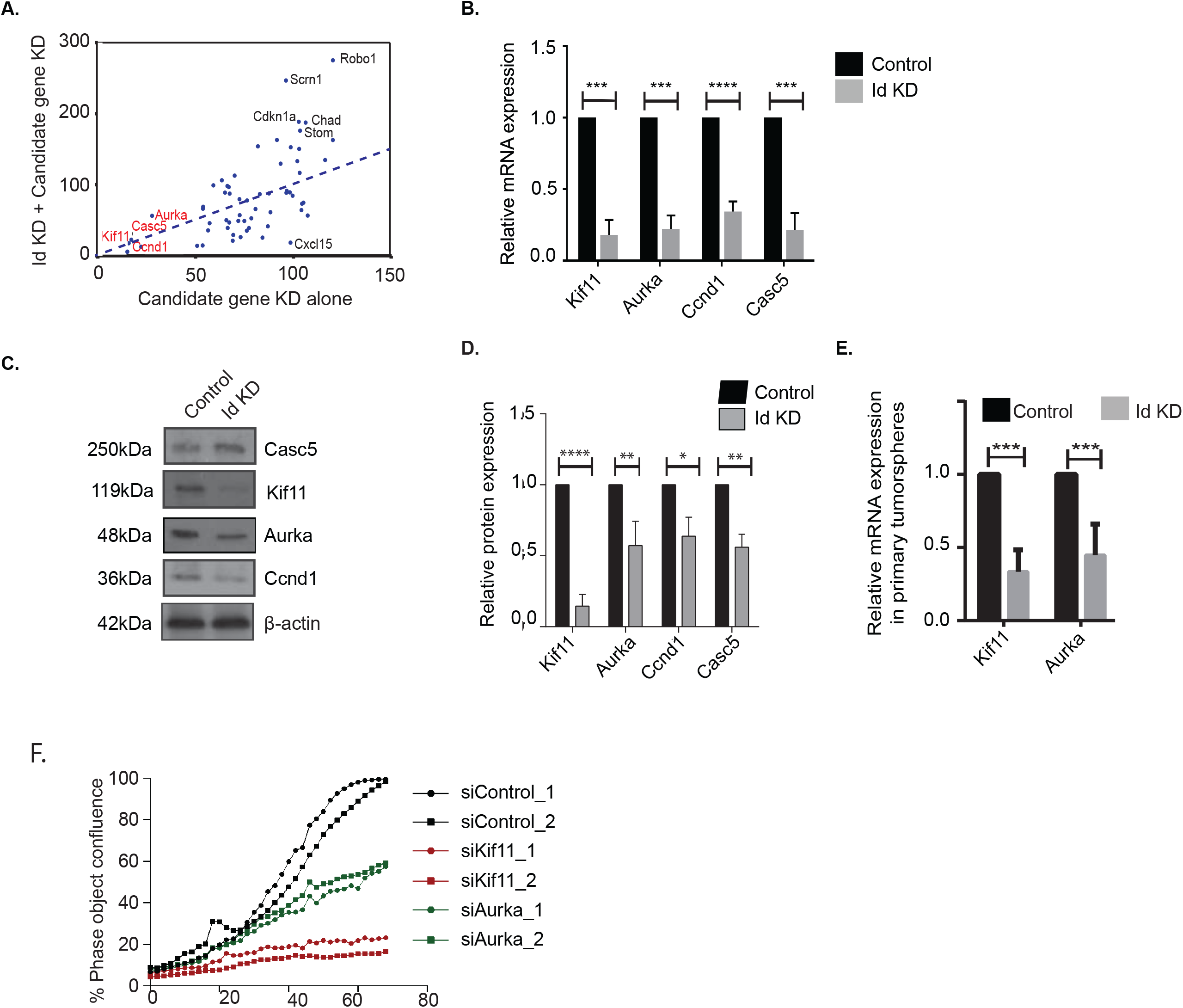
Effect of target gene knockdown on the proliferative phenotype. A) The effect of the candidate genes on proliferation of4T1 cells was asseses by reverse transfection with 40nM siGENOMESMART pool siRNA against each of the 57 candidate genes. Cell viability was quantified at 72h post-transfection using the CellTiter-Glo® luminescent assay. IncuCyteZOOM® live cell imaging every 2 hours was also performed, which allowed us to quantify cell growth (confluence) over time throughout the experiment. Interestingly, the target genes that showed the greatest effect on the viability and thus the proliferative phenotype of the 4T1 cells with more than 50% were Kif11,Ccnd1, Casc5, and Aurka. (B) The relative mRNA expression level of Kif11, Aurka, Ccnd1 and Casc5 in Id KD cells with respect to control cells were quantified with qRT-PCR. Data were normalized to beta-actin and analyzed by the 2−ΔΔCt method. (C) The expression of main target genes Casc5, Kif11, Aurka, Ccnd1 at protein level were analysed in both control and Id KD cells by western blot.(D) Quantification of relative protein expression of Kif11, Aurka, Ccnd1 and Casc5 in control and Id KD, normalized with b-actin showed a decrease in knock down conditions. (E) Relative mRNA expression level of Kif11, Aurka in primary tumour spheres was assesed. (F) Phase object confluency on Kif11 KD and Aurka KD showing significant decrease in cell proliferation compared to the Controls.

To investigate the effect of putative Id targets such as Kif11, Aurka, Casc5 and Ccnd1 on cell cycle, relative mRNA levels were analysed. We observed significant reduction in the mRNA levels of Kif11, Aurka, Ccnd1 and Casc5 in Id KD compared to the controls (Figure 3B). We also detected a decrease in the expression of Kif11, Aurka, Ccnd1 and Casc5 at the protein level by western blot (Figure 3C, D). Kif11 and Aurka were also down regulated at the transcript level in spheres generated in the Id KD cells when compared to control (Figure 3E, Supplementary Figure 2D).

To confirm the effect of putative Id targets on proliferation, we used an independent pMission siRNA system in 4T1 parental cells. Loss of Kif11 and Aurka lead to significant decrease in the proliferative capacity of the 4T1 cells (Figure 3F, Supplementary Figure 3A, B). We continued our studies with Kif11 and Aurka as we did not observe any significant loss of proliferative phenotype with Casc5 and Ccnd1 knock down (Supplementary Figure 3C, D).

### The effect of Id KD on key molecular cell cycle genes is different from Kif11 and Aurka

As proliferation is inextricably linked to the cell cycle, we next characterized the effect of Kif11 and Aurka knock down using the pMission system on the cell cycle. We found that Kif11 and Aurka depletion lead to cell cycle arrest in the G2/M phase as evidenced by cell cycle analysis (Figure 4 A,B,C). Intriguingly, this observation was fundamentally different from the G0/ G1 arrest that cells undergo when Id is depleted (Figure 1A, Supplementary 3E).

**Figure 4.**
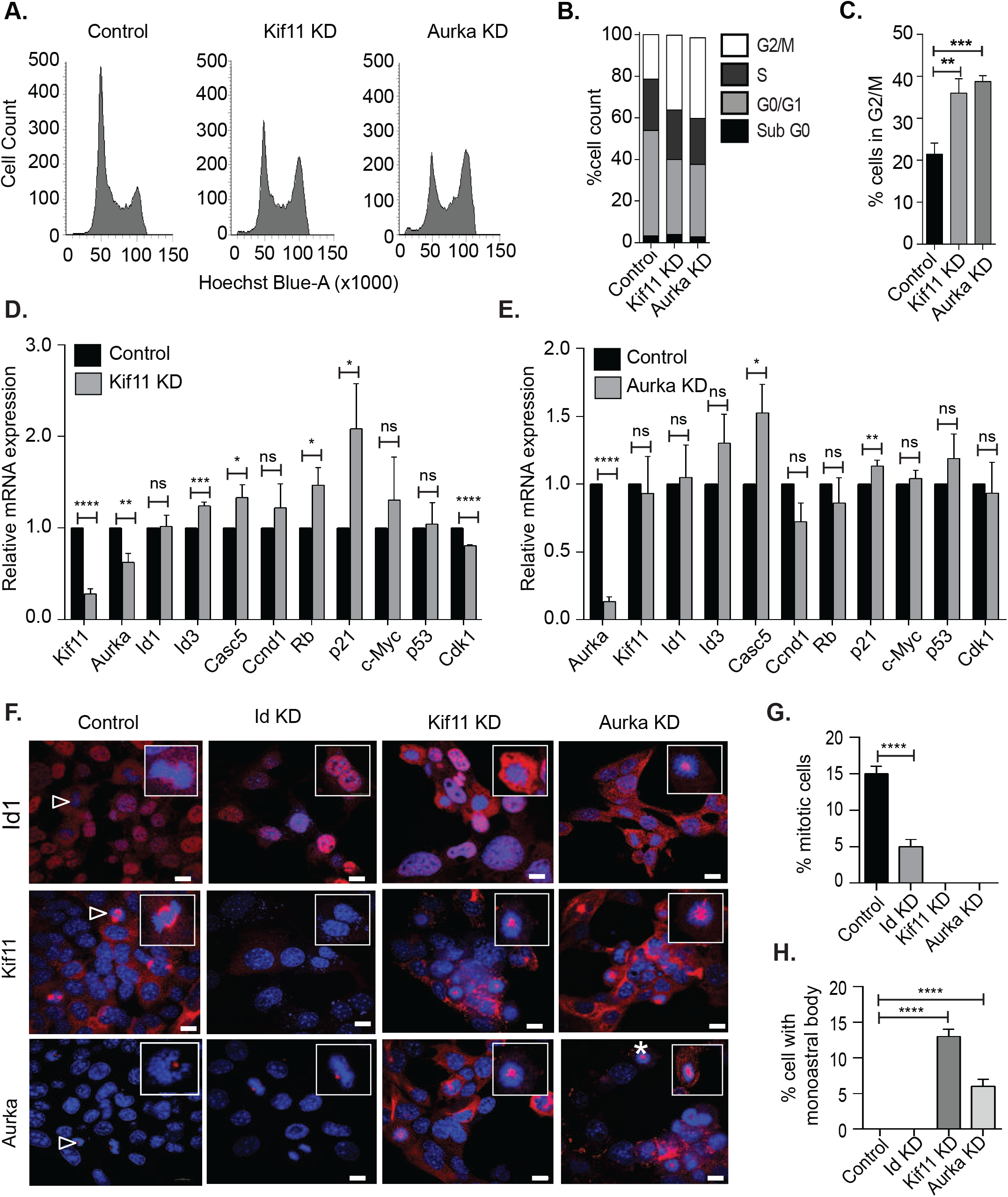
Kif11 and Aurka depletion does not phenocopy loss of Id1. (A) Flow cytometric analysis of cells after knocking down Kif11 (Kif11 KD), Aurka (Aurka KD) along with scrambled control (Control). (B) Comparing the percentage cell count in each phase of the cell cycle after knocking down Kif11 or Aurka. (C) Percentage of cells in G2/M phase in Control, Kif11 KD, Aurka KD. Statistical significance was analyzed by an unpaired t-test. ***p< 0.001, **p< 0.01, *p< 0.05. (D)The relative mRNA expression level of important cell cycle genes were analysed in Kif11 KD and Aurka KD (E) with respect to scrambled control using qRT-PCR. Data were normalized to beta-actin and analyzed by the 2−ΔΔCt method. Statistical significance was analyzed by an unpaired t-test. ***p< 0.001, **p< 0.01, *p< 0.05. (F) shows the Immunofluoresence staining for Id1, Kif11 and Aurka on 4T1 control, Id KD, Kif11 KD andAurka KD cell lines. Representative images are taken using Nicon A1R+ confocal system at 100x magnification with scale bar corresponds to 100um. Dapi shows the nuclear staining, aster(*) shows the monoastral spindle formation in siRNA KD system, Δ shows normal cell division, inset shows the 100x zoomed images of the same. (G) shows the percentage of cells in M-phase in Control, Id KD, Kif11 KD and Aurka KD cell lines. (H) Quantification of cells exhibiting monoastral bodies in Control, Id KD, Kif11KD and Aurka KD conditions.

To determine the molecular effect of Kif11 and Aurka on cell cycle, relative mRNA levels of key cell cycle genes were analysed. Kif 11 KD significantly reduced the expression of Aurka even though the expression of Id1 was not altered (Figure 4D). Decreased Kif11 expression had a positive effect on the mRNA levels of Id3, Casc5, Rb and p21. It reduced the expression of Cdk1, but had no significant effect on Ccnd1, c-Myc and p53. Knock down of Aurka did not show any effect on Id1, Id3 and Kif11, but decreased the mRNA level of Ccnd1 and increased that of Casc5 and p21 (Figure 4E). Altogether these results suggests that Id is having an upstream effect on Kif11 and Aurka is possibly a downstream target of Kif11.It also gives a hint that Id is acting through these target molecules via a unique molecular pathway which is independent of the cell cycle.

### Kif11 or Aurka depletion does not phenocopy loss of Id

We next compared our microscopic observations on the phenotype of the Id system with that of depleting the cells of the Id putative targets, Kif11 and Aurka. We noticed monoastral bodies with misformed mitotic spindle, indicating that the majority of the Kif11 KD and Aurka KD cells are arrested in M phase of the cell cycle (Figure 4F, Supplementary Figure 3E)). The formation of monoastral bodies is indicative of duplicated chromatin (4N) not being able to separate due perturbations in the spindle and centrosome, thus indicates G2/M arrest and matches the cell cycle analysis (22) (Figure 4A). Quantification of cells in M-phase (Figure 4G) and cells exhibiting the monoastral body phenotype (Figure 4H) in Control, Id KD, Kif11 KD, and Aurka KD clearly demonstrates that Id depletion results in a phenotype that is distinct from Kif11 and Aurka which points to a unique mechanism controlled by Id.

### Therapeutic targeting of CSCs through Id1-Kif11/Aurka axis

There is currently no effective targeted therapy for TNBC and chemotherapy is usually the first line of treatment with a relapse rate of 25% (23). Our previous work has demonstrated that Id is critical for CSC associated phenotypes in TNBC such as proliferation, self-renewal, migration and metastasis(1). We have now identified that these phenotypes are controlled by the Id-Kif11/Aurka axis. We hypothesized that targeting Kif11 or Aurka to block the Id1-Kif11/Aurka axis may make the Id expressing CSC more vulnerable to chemotherapy, thus more effectively debulking the entire tumor mass.

To test this hypothesis, we first determined the IC50 values for two commonly used chemotherapy drugs in breast cancer treatment, Paclitaxel and Doxorubicin, in 4T1 cells (Supplementary Figure4A). Interestingly, we find that there is a significant enrichment for Id1+ tumor cells after treatment with Doxorubicin and Paclitaxel, suggesting that the Id1+ CSCs are chemo resistant (Figure 5A). We next determined the IC50 values for the small molecule inhibitors of Kif11 and Aurka, Ispinesib and Alisertib, respectively (Supplementary Figure 4B). Cells treated with Ispinesib showed a significant reduction in the percentage of Id1+ cells but no significant change was observed in those treated with Alisertib as compared to the Control (Figure 5A). We decided to continue with Paclitaxel and Ispinesib based on these results.

**Figure 5.**
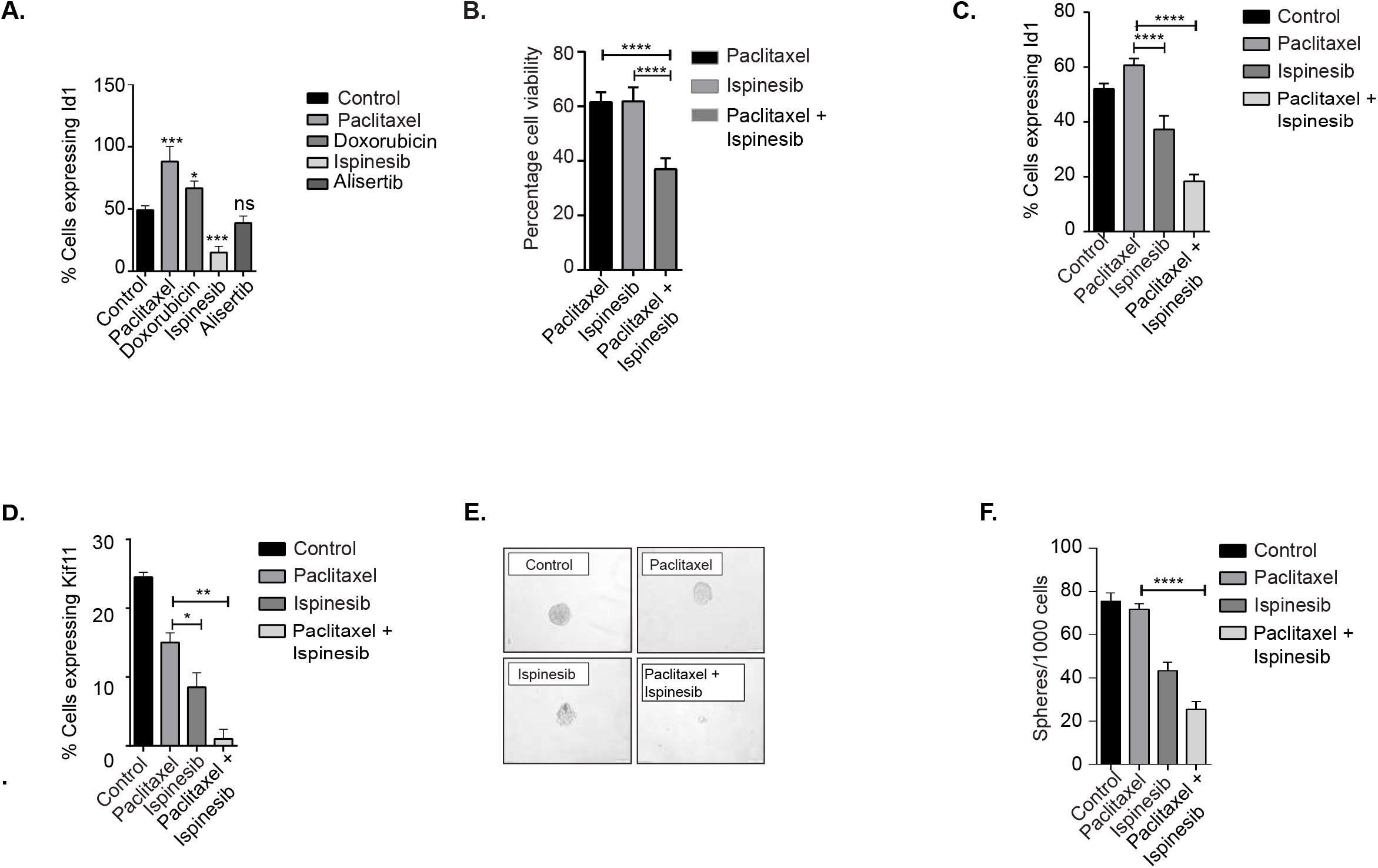
Therapeutic targeting of CSC through Id1-Kif11/Aurka axis. (A) Cells expressing Id1 after treatment with Paclitaxel, Doxorubicin, Ispinesib and Alisertib compared to the Control. (B) Cell viability was determined using the MTT assay after treating the cells with Paclitaxel, Ispinesib and the combination therapy of Paclitaxel + Ispinesib. (C) Percentage cells expressing Id1 in Control, Paclitaxel, Ispinesib and combination treatment showing a significant decrease in Id1 cells when treated with Ispinesib and combination of chemotherapy and small molecule inhibitor. (D) Percentage cells expressing Kif11 under control, Paclitaxel, Ispinesib and combination treatment. (E) Phase contrast image showing the tumour sphere size under Control, Paclitaxel, Ispinesib and combination treatment (F) There is a significant reduction in sphere forming ability with combination treatment when compared to chemotherapy Paclitaxel alone.

We next asked whether the reduction in Id1 levels by Ispinesib can increase the sensitivity of these cells to Paclitaxel in TNBC cells. As proof of principle, we found that the survival fraction and Id1, Kif11 expression in cells treated with a combination of Paclitaxel and Ispinesib was significantly less when compared to cells treated with either Paclitaxel or Ispinesib alone (Figure 5B,C, D). This suggests that the disruption of the Id1-Kif11 axis by Ispinesib sensitizes CSCs to chemotherapy.

We next checked the effect of combination therapy on *in vitro* self-renewal of TNBC cells. A significant reduction in the self-renewal capacity was observed in cells treated with the combination of Paclitaxel and Ispinesib when compared to cells treated with either of the drugs alone (Figure 5 E,F). The expression levels of Id1and Aurka decreased upon combination treatment when compared to the chemotherapy Paclitaxel (Supplementary Figure 4C). We have also checked the expression of known CSC markers CD24/CD29 in these cells and interestingly found that the CSC fraction was significantly reduced with combination therapy when compared to using Paclitaxel alone (Supplementary Figure 4D).

In conclusion we show that targeting the Id1/ Kif11 molecular pathway in the Id1+ CSCs in combination with chemotherapy results in more effective debulking in TNBC.

## Discussion

The current body of work sheds light on the role Id proteins (specifically Id1 and Id3) play in affecting the CSC phenotype of proliferation due to the striking G0/G1 cell cycle arrest we observed when cancer cells were depleted of Id proteins. The Gap phase (G1) is not simply a time delay between the M and S phase. It is a time period within which the cell can monitor the internal and external environment to ensure that the conditions are optimal for the S and M phases. The G1 phase is especially important as the cell can stop in a special G0 resting phase if it finds the conditions are unfavourable for the cell to undergo further cell division. The most distinctive feature of Id depleted cells is the lack of DNA division as also reflected in the G1 block. Combining the phenotype with the change in expression levels of critical cell cycle genes, Id pauses or checks the cells in the G1 state in a manner that they can re-enter the normal cell cycle once the stress is removed. This supports the theory of Id as a master regulator that on sensing unfavourable conditions, can “brake” the cells in the G1 phase through multiple means (molecular regulation of cell cycle genes, DNA division inhibition, protein complex perturbation at the centrosome and spindle fibres). This strategy would allow cells to survive in a state of stasis till conditions favourable to growth of the tumor cell arise. Interestingly, this state could be reversed suggesting that the cells are poised in a state of cell arrest under unfavourable conditions (like chemotherapy treatment) and re enter the cell cycle subsequently.

The idea that CSCs are more plastic and can exist in more than one state may be supported by looking at the EMT program. This makes sense from the point of view of the EMT scores, i.e. the Id KD is not pushing the cells clearly towards a more E or a more M state, as the levels of both canonical markers decrease. Also, our bioinformatic model uses a ratio of CDH1/VIM as a predictor to calculate the scores; so the relatively proportional change that we see at RNA/protein levels for both CDH1 and VIM is consistent. The data adds to the evidence that EMT and MET are not binary (24)for different stages of EMT and their varying degrees of causal contribution to metastasis.

Using two independent models of Id1 gene expression and gene depletion, we were intrigued to identify the critical cell cycle genes, Kif11and Aurka as Id putative gene targets. Interestingly previous work in nasopharyngeal cells have linked Id1 and Aurka mechanistically in the induction of tetraploidy. Id1 was found to affect Aurka degradation which normally occurs during exit from mitosis by the APC/C Cdh1 mediated proteolysis pathway. Id1 stabilized Aurka by actively competition with Cdh1, thus preventing Aurka degradation(25). Interestingly, while individual knock down of Kif11 and Aurka also led to a proliferative arrest, it did not phenocopy the G0/G1 cell cycle arrest with the Id knock down or the formation of monoastral bodies. This suggested that the impediment of the cell cycle by Id protein is through different mechanisms and not the canonical mitotic pathways involving the microtubules by Kif11/ Aurka which forms a part of our future investigation.

There is no targeted therapy for TNBC and chemotherapy is the first line of treatment. Thus, we checked the effect of commonly used chemotherapy drugs paclitaxel and doxorubicin which are used in the clinic for TNBCs. Studies by our group and others have already reported that Id1 marks a chemoresistant breast cancer cell in cancers like Hepatocarcinomas. Interestingly, previous work has targeted the Kif11 pathway in Docetaxel resistant TNBC cells(26). But the most compelling reason to target the Id1/ Kif11 pathway came from work by Chattopadhya et al(27) who identified the drug BRD9876 as a Kinesin-5 inhibitor in Multiple Myeloma which led to significant down regulation of ID1. Based on our work, we used Id as a marker for the chemoresistant CSC population in TNBC. We tested our hypothesis that we can achieve a better response by combining traditional chemotherapy along with ablation of the Id expressing chemoresistant cells using small molecule inhibitors against the Id targets Kif11. We achieved a significant decrease in proliferative and self renewal capacity when the cells were treated with Paclitaxel and Ispiniseb by successfully targeting sub populations of cells including the Id+ CSCs within a tumor.

Thus a combination of targeted drugs with chemotherapy would be an effective strategy for the complete treatment of TNBC and give women currently living with this disease a better long term prognosis.

## Materials and Methods

### Mammalian cell culture

The cell lines 4T1pSLIK cell lines and Parental cell lines used in this study were obtained from the American Type Culture Collection (ATCC). All cell lines were cultured at 37°C in 5% CO2 and 95% air to no more than 80% confluence. Cells were passaged by washing with PBS twice and trypsinised with trypsin-EDTA (0.05%), followed by incubation at 37°C until the cells were detached from the tissue culture flask. An equal or greater volume of culture medium was added to neutralise the trypsin-EDTA. Appropriate volume of cell suspension was then added into a new tissue culture flask for passaging. All cell lines were preserved by cryopreservation. Each cryovial contained 1×106 cells and were frozen in a solution consisting of 50% (v/v) foetal bovine serum (FBS), 40% (v/v) growth media, and 10% (v/v) DMSO to −80°C at a rate of 1°C/min for a minimum of 4hr before transferring to liquid nitrogen. Cells were revived by warming individual cryovials to 37°C and seeding into 10mL of culture medium in a T75 tissue culture flask.

### Immunofluoresence

Cells were seeded on coverslips in tissue culture dishes and cultured for 2 days. Cells were washed in PBS and fixed with 4% paraformaldehyde. Fixed cells were washed with PBS and resuspended in 0.2% Triton-X (Sigma) in PBS solution for 20 minutes. Cells were blocked using 1%BSA in PBS for 1 hr at room temperature and primary antibody was added and incubated at 4 degrees overnight. Next day the cells were washed with PBS thriceand secondary antibody was added with Dapi(1mg/ml)and incubated for 1 hr at room temperature. Cover slips were mounted in Prolong gold antifade reagent and visualized under Fluorescence microscope.

### Microscopic imaging

Cells on tissue culture plates were magnified with Fluorescence microscope (Olympus, Germany), under both high and low magnification. Confocal images were captured by the Leica DFC280 digital camera system (Leica Microsystems, Wetzlar, Germany).

### Cell Cycle Analysis

Cells were harvested and spin down at 1200rpm for 5 minutes. Cells were counted and Hoechst (Sigma) (4ug/mL) was added to the cell suspension(1 million cells) and incubated at 37^0^C for 30 minutes. The cell cycle distribution was determined with a flow cytometer (BD Aria III).The data were analysed using the BD FACS analyser software.

### RNA extraction and Real-time PCR (RT-PCR)

Total RNA was isolated using TRIzol® Reagent (Invitrogen) and cDNA was synthesized using High-Capacity cDNA Reverse Transcription Kits (Applied Biosystems; Thermo fisher Scientific, Inc.). Real-time PCR was performed on the QuantStudio 7 Flex Real-Time PCR System using Power SYBR Green PCR Master Mix (Applied Biosystems). The PCR conditions were 950C for 10 min, followed by 40 cycles of 950C for 30s, and 600C for 1 min. All reactions were done in triplicates and the transcript levels were normalized to those of b-act. The relative fold change was determined by 2−ΔΔCT method as described (PMID: 11846609). The gene specific primers used for RT-PCR are listed in (Table)

### CSC markers staining

The cells were collected at 1200rpm for 5 minutes at 4°C(There should be atleast 1 million cells). Pellets were resuspended in FcBlock (Miltenyibiotec, 1:10) and incubated on ice for 10minutes. Cells were pelleted (1200rpm, 5mins, 4°C) and washed by resuspending with cold PBS+salts, then pelleted again. Cells were resuspended in lineage marker cocktail CD29 (Miltenyibiotec, 1:10), CD61 (Miltenyibiotec, 1:10) and incubated on ice for 20 minutes. Cells were pelleted, washed with PBS and pelleted. Cells were resuspended in FACS buffer1:400) and incubated on ice for 20 minutes. Cells were pelleted, washed with PBS+salts, pelleted then resuspended in 200ul FACS buffer(PBS containing salt + 2% FBS + 2% HEPES). Appropriate single stain and unstained controls were performed alongside CSC marker staining.

### Microarray and bioinformatics analysis

Total RNA from the samples were isolated using Qiagen RNeasy minikit (Qiagen, Doncaster, VIC, Australia. cDNA synthesis, probe labelling, hybridization, scanning and data processing were all conducted by the Ramaciotti Centre for Gene Function Analysis (The University of New South Wales). Gene expression profiling was performed using the AffymetrixGeneChip® Gene 1.0 ST Array, a whole-transcript array which covers >28000 coding transcripts and >7000 non-coding long intergenic non-coding transcripts. Data analysis was performed using the Genepattern software package from the Broad Institute. Three different modules, Hierarchical Clustering Viewer, Comparative Marker Selection Viewer and Heatmap Viewer were used to visualize the data. In addition to identifying candidate molecules and pathways of interest, Gene Set Enrichment Analysis (GSEA) (http://www.broadinstitute.org/gsea) was performed using the GSEA Pre-ranked module. Briefly, GSEA compares differentially regulated genes in an expression profiling dataset with curated and experimentally determined sets of genes in the MSigDB database to determine if certain sets of genes are statistically over-represented in the expression profiling data

### siRNA screen to assess proliferation

Reverse transfection of 4T1 cells in 384 well plates was performed with 400 cells and 0.08uL Dharmafect1 per well using a Caliper Zephyr and Biotek EL406 liquid handling robots. Media was change at 24hr post-transfection. Cell titerglo assay was performed using a BMG Clariostar plate reader (luminescence assay). Final data presented is generated from three biological replicates each consisting of two technical replicates. Viability measurements were normalized to the treatment-matched scrambled control after subtracting the blank empty wells.

4T1 cells were reverse transfected with a 40nM siGENOMESMART pool siRNA against each of the 57 candidate genes. Cell viability was quantified at 72h post-transfection using the CellTiter-Glo® luminescent assay (Table 1). IncuCyte ZOOM® live cell imaging every 2 hours was also performed, which allowed us to quantify cell growth (confluence) over time throughout the experiment.

### Extracting protein lysates

Protein lysates were obtained by direct lysis from the tissue culture plates. Cells were washed once with chilled PBS (Invitrogen, India) before adding ice cold RIPA lysis buffer supplemented with complete protease inhibitor cocktail solution (Sigma Aldrich, India) to inhibit protein degradation. The cell lysate was then transferred to 1.5mL tubes. All steps above were performed on ice. The cell lysates were then centrifuged at 14000rpm for 10min, 4°C. The supernatants were transferred to new 1.5mL tubes and were stored at −80°C for later use or on ice for immediate use.

### Quantifying protein concentration

The protein concentration of each sample was measured by using a BCA assay using the micro bicinchoninic acid (BCA) kit (Thermofisher scientific, India) following the manufacturer’s instructions. The assay was performed in a clear-bottomed flat surface 96-well plate. Briefly, the BSA (2mg/ml) was serially diluted in distilled water to generate a dilution range of: 0.0μg/μL to 2μg/μL. Protein lysate was diluted 1 in 10 with distilled water. The BCA reagents were then mixed (50:1 Part A:Part B) and 200ul was added to each well. The plate was then incubated at 37°C for 30min, followed by a measurement of absorbance at 562nm using the TECAN plate reader. The protein concentration of each sample was calculated by using GraphPad prism software.

### Sodium dodecyl sulphate polyacrylamide gel electrophoresis (SDS-PAGE)

SDS was performed by using the Biorad system (Biorad, India). 20μg of protein from each sample was made up in 1x Lammelli sample buffer and denatured at 95°C for 10min. The denatured protein samples were loaded onto polyacrylamide gels. Gels consisted of a 5% acrylamide stacking gel and a 12% acrylamide gradient separating gel. Electrophoresis was performed for 45min at 140V in 1x SDS running buffer.

### Protein transfer and immunoblotting

Following electrophoresis, proteins were transferred onto PVDF membranes (Biorad, India) at 120V for 50min in 1x transfer buffer (Biorad, India). The PVDF membranes were blocked in a solution containing 5% (w/v) skim milk powder and 0.1% TBS-tween for 1hr at room temperature. After blocking, the membranes were washed 3 times in TBS-tween (5min each time). The primary antibodies were diluted in an antibody diluting solution containing 5% (w/v) BSA, 0.025% (w/v) sodium azide in 1% (v/v) TBST. The washed membranes were incubated with primary antibody solutions at concentration as per Table for 1hr at room temperature or overnight at 4°C. Following primary antibody incubation, the membranes were washed 3 times (10min each time). The secondary antibodies used were anti-rabbit IgG or anti-mouse IgG conjugated with horseradish peroxidise (HRP). Membranes were incubated in secondary antibodies at a concentration of 1:10000 in 5% (w/v) skim milk/TBS-tween buffer for 1hr at room temperature. Excess secondary antibody was washed in TBS-tween four times (15min each time). Specific protein bands were detected by ECL chemiluminesence (Biorad, India).

### MTT assay

4 T1 cells were seeded at a density of 500 cells/well in 96 well plate.When the cells become 80% confluency, freshly prepared MTT reagent (5ug/mL) was added to the culture media. Incubate the plate for 3 hrs in dark at 37^0^C. Remove the media containing reagent from each well and add 200ul of DMSO to each well. Take the absorbance readings using a microplate reader at 570nm.

### IC50 values for Chemotherapeutic drugs

4T1 cells were harvested and seeded 1000 cells/well in 96 well plate. When the cells become 20% confluency, culture media was removed and replenished with media containing chemotherapeutic drugs. MTT reading was taken 48hrs post drug treatment.

### Tumoursphere assay

4T1 cells were trypsinised and washed twice with PBS (Invitrogen, India). Cells were then resuspended in RPMI1640 medium without FBS and sieved through a 40μM cell strainer (BD Falcon, India) twice to ensure at least 95-99% of cells were in single cell suspension before being counted on the haemocytometer. Single cells were plated in ultralow attachment 6-well plates (Corning, India) at a density of 2.0 x 104 viable cells/well in triplicate. Cells were cultured in serum-free RPMI1640 medium, supplemented with B27 (Invitrogen, India) and 20ng/mL bFGF (Millipore, India) and 4μg/mL heparin (Sigma-Aldrich, India). Serum-free media supplemented with the additives mentioned above was added to the cells every 3 days. The plate was tapped very gently to ensure even distribution of the cells. Primary tumourspheres were counted at day 8.

### Statistical analysis

Statistical analyses of the data were performed using GraphPad Prism 6. All in vitro experiments were done in 3 biological replicates each with 2 or more technical replicates. Data represented are means ± standard deviation. Statistical tests used are Unpaired student t-test and two-way-ANOVA. p-values<0.05 were considered statistically significant with *p<0.05, **p<0.01, ***p<0.001, **** p< 0.0001.

## Declarations

### Consent for publication

All authors have given their consent for publication.

### Author’s contributions

RN contributed to the conceptualizaton. RS, BAV, NK and APT contributed to the methodology. DLR to the genomic analysis. RM, BAV, APT, NK, AC, ED, AM, CK and HH contributed to the investigations. RN wrote the original draft of the manuscript. JTG, HL and MKJ contributed to the bioinformatic analysis of the EMT program. AS reviewed and edited the manuscript. AS and RN contributed to the funding acquisition. All authors read and approved the final manuscript.

## Acknowledgements

We would like to thank the following people for their assistance with this manuscript: Mr. Tilak Prasad and Mrs. Surabhi S.V for helping with Flow cytometry. Mr. Anurup K.G for technical support with Confocal imaging. Dr. Vishnu Sunil Jayakumar for Animal Handling. Dr. T. R. Santhosh Kumar’s Lab for helping with lab consumables and technical support.

## Author information

Radhika is senior author.

---

**Supplementary Figure 1.**
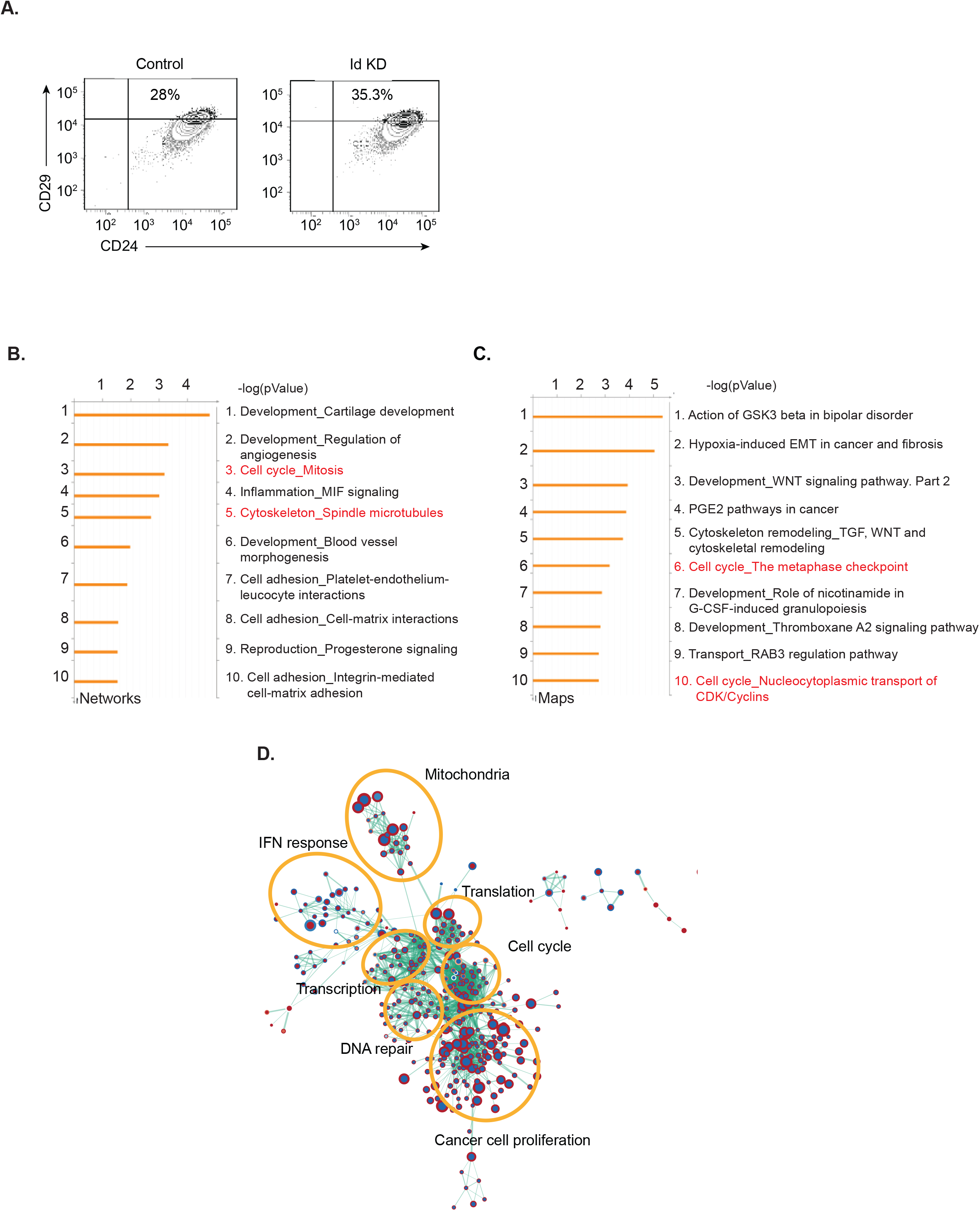
Identification of putative ID regulated genes. (A) Cancer stem cell markers CD29/CD24 in Id KD conditions is not significantly changed from controls. (B) The gene expression profile of three independent replicates (with and without doxycycline treatment), was compared by microarray analysis to generate a list of differentially expressed genes between Id depleted and control 4T1 cells. (C) The gene expression profiles of the Id+ and Id-cells from three independent Id1C3Ttg tumours were compared by RNA sequencing. This resulted in a list of differentially expressed genes between the Id+ and Id-mouse TNBC tumour cells. (D) Aiming to discover high confidence genes involved in the Id gene regulatory network, lists of differentially expressed genes between the Id expressing or depleted TNBC models and their controls were compared using MetaCoreTM software. By comparing these two datasets, lists of MetaCore network objects common to both experiments as well as those unique to each of the two data sets was generated. Results are visualized using the enrichment map plug-in for Cytoscape. Each circular node is a gene set with diameter proportional to the number of genes. The outer node color represents the magnitude and direction of enrichment (see scale) in Id1C3Tag cells, inner node color enrichment in Id KD cells. Thickness of the edges (green lines) is proportional to the similarity of gene sets between linked nodes. The most related clusters are placed nearest to each other. The functions of prominent clusters are shown.

**Supplementary Figure 2.**
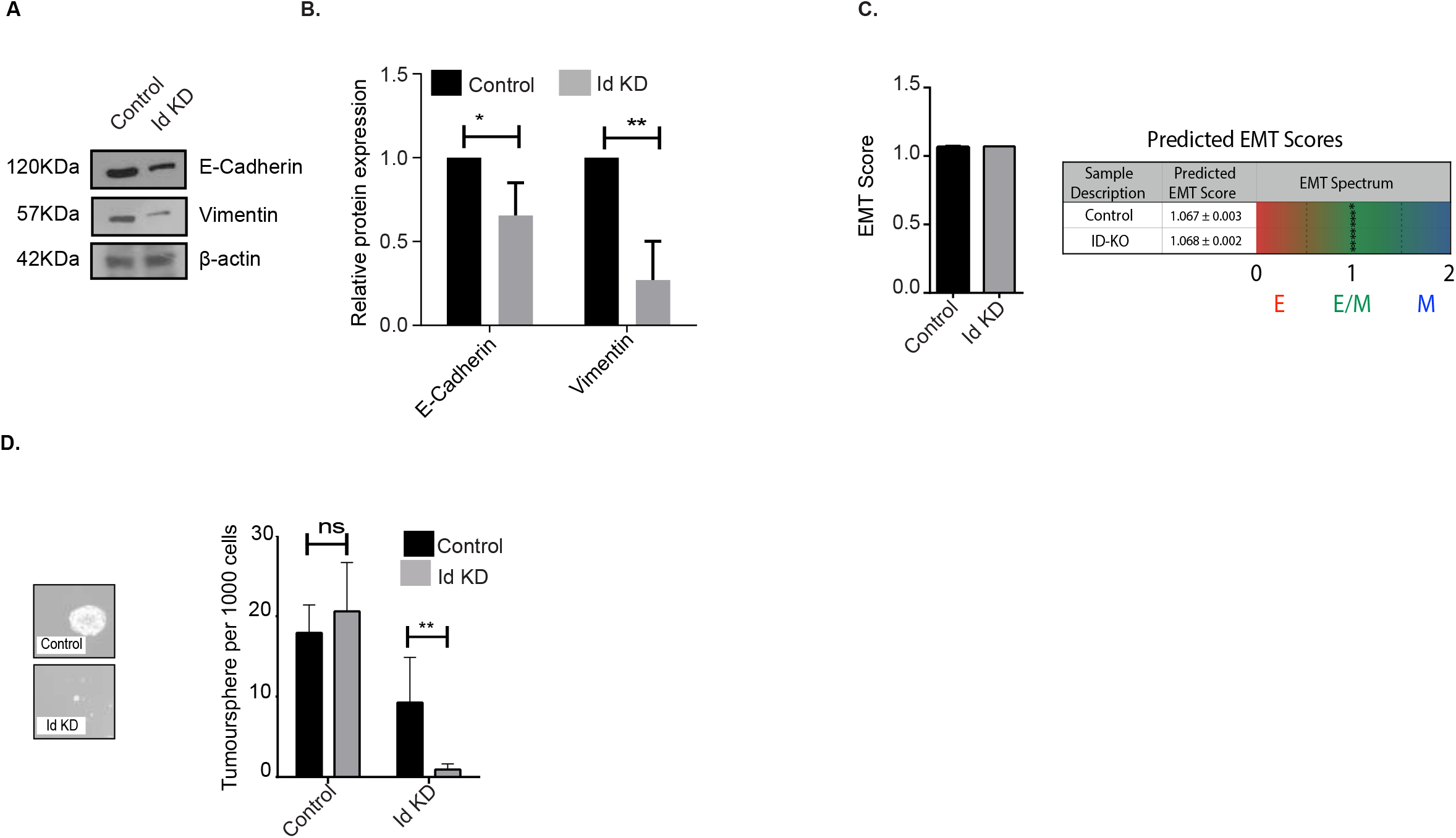
Effect of Id knockdown on Id targeted genes. (A) and (B) Relative protein expression level and quantification of E-cadherin, Vimentin in Id KD cells with respect to Control cells were quantified with western blot. β-actin was used as the loading control. (C) EMT score calculated for all samples, on a scale of 0 (fully epithelial) to 2 (fully mesenchymal). (D) Phase contrast images and quantification of primary spheres under control and Id KD conditions.

**Supplementary Figure 3.**
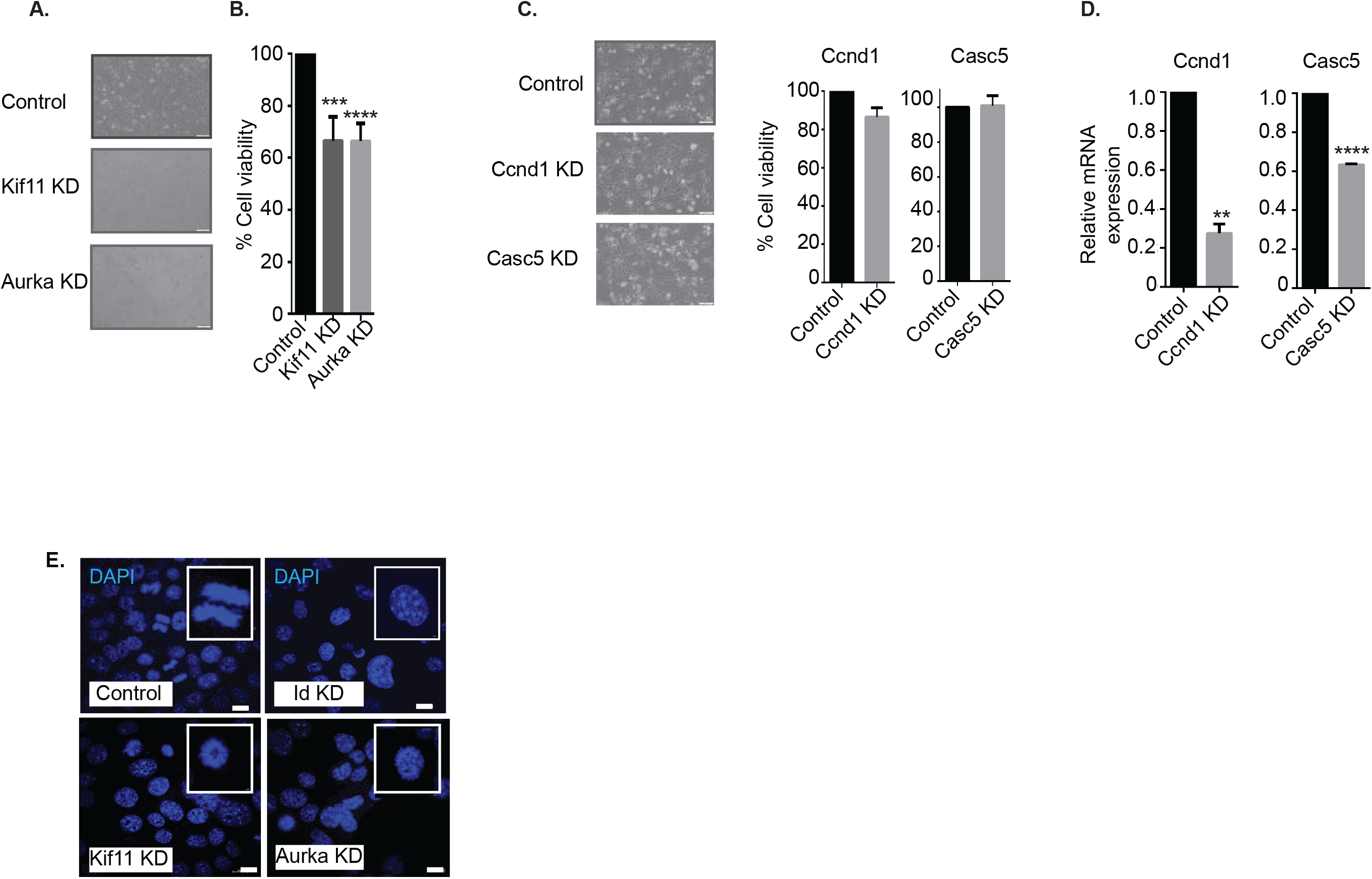
Id KD leads to cell cycle arrest. (A) Phase contrast images of 4T1 cells under control, Kif11 KD and Aurka KD. (B) A significant decrease in cell viability was determined in Kif11 KD and Aurka KD cells compared to Controls. (C) Phase contrast images and cell viability of 4T1 cells under Control, Ccnd1KD and Casc5 KD conditions.(D) Relative mRNA expression level of Ccnd1 and Casc5 under Ccnd1 and Casc5 KD condition compared to the control. (E) Immunofluorescence images showing Dapi staining of Control, Id KD, Kif11 KD and Aurka KD cells under fluorescence microscope. Representative images are taken using Nikon A1R+ confocal system at 100x magnification with scale bar corresponds to 100um, inset shows the 100x zoomed images of the same.

**Supplementary Figure 4.**
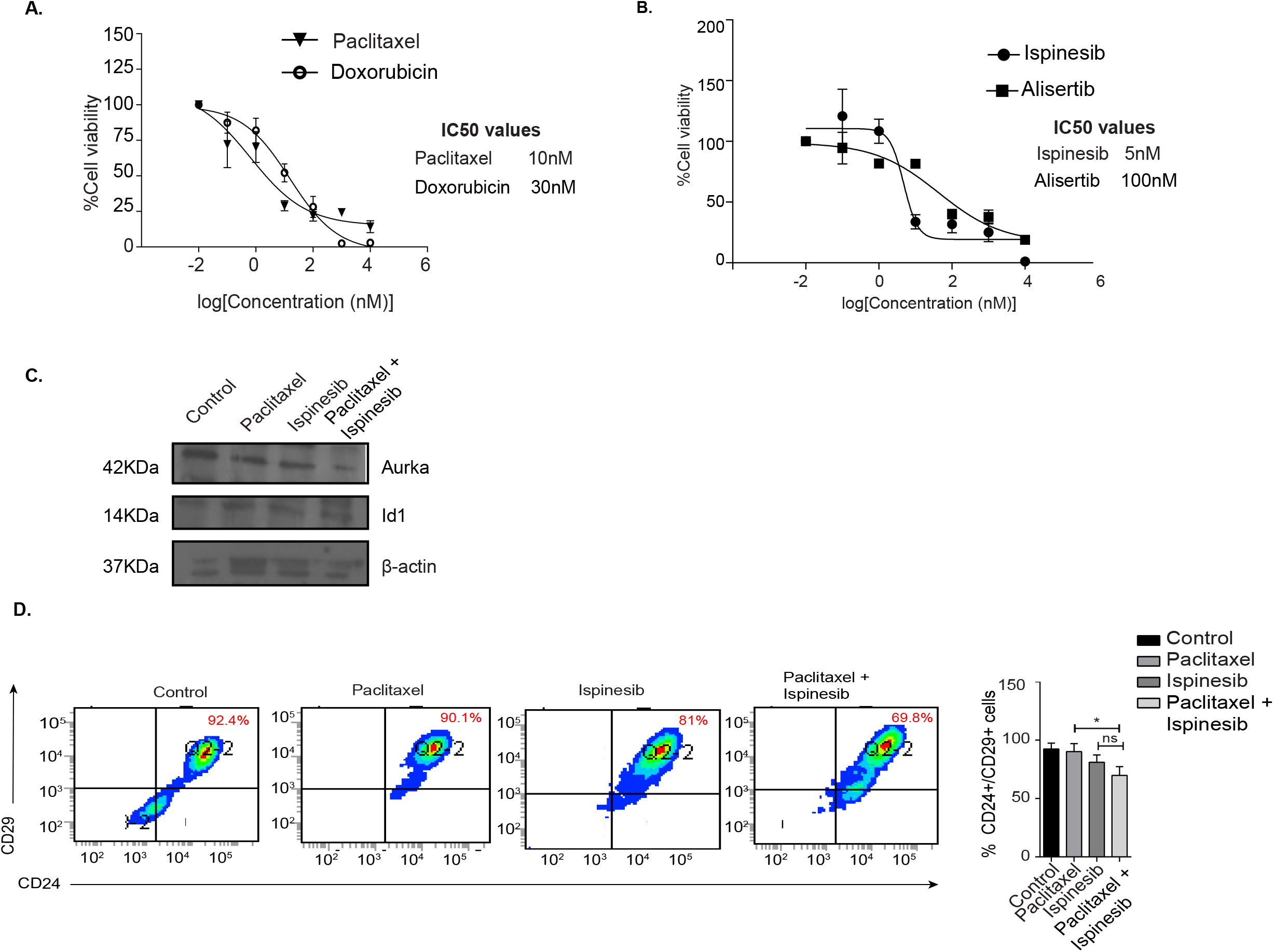
Combination therapy is more effective. (A) IC50 values in 4T1 cells for two commonly used chemotherapy drugs in breast cancer treatment, Paclitaxel and Doxorubicin. (B) IC50 values for the small molecule inhibitors of Kif11 and Aurka, Ispinesib and Alisertib. (C) Relative protein level expression of Id1, Kif11 and Aurka on Paclitaxel, Ispinesib and Paclitaxel +Ispinesib treated 4T1 cells. (D) Expression of known CSC markers CD24/CD29 on single and combination treated 4T1 cells.

**Supplementary Table 1.**
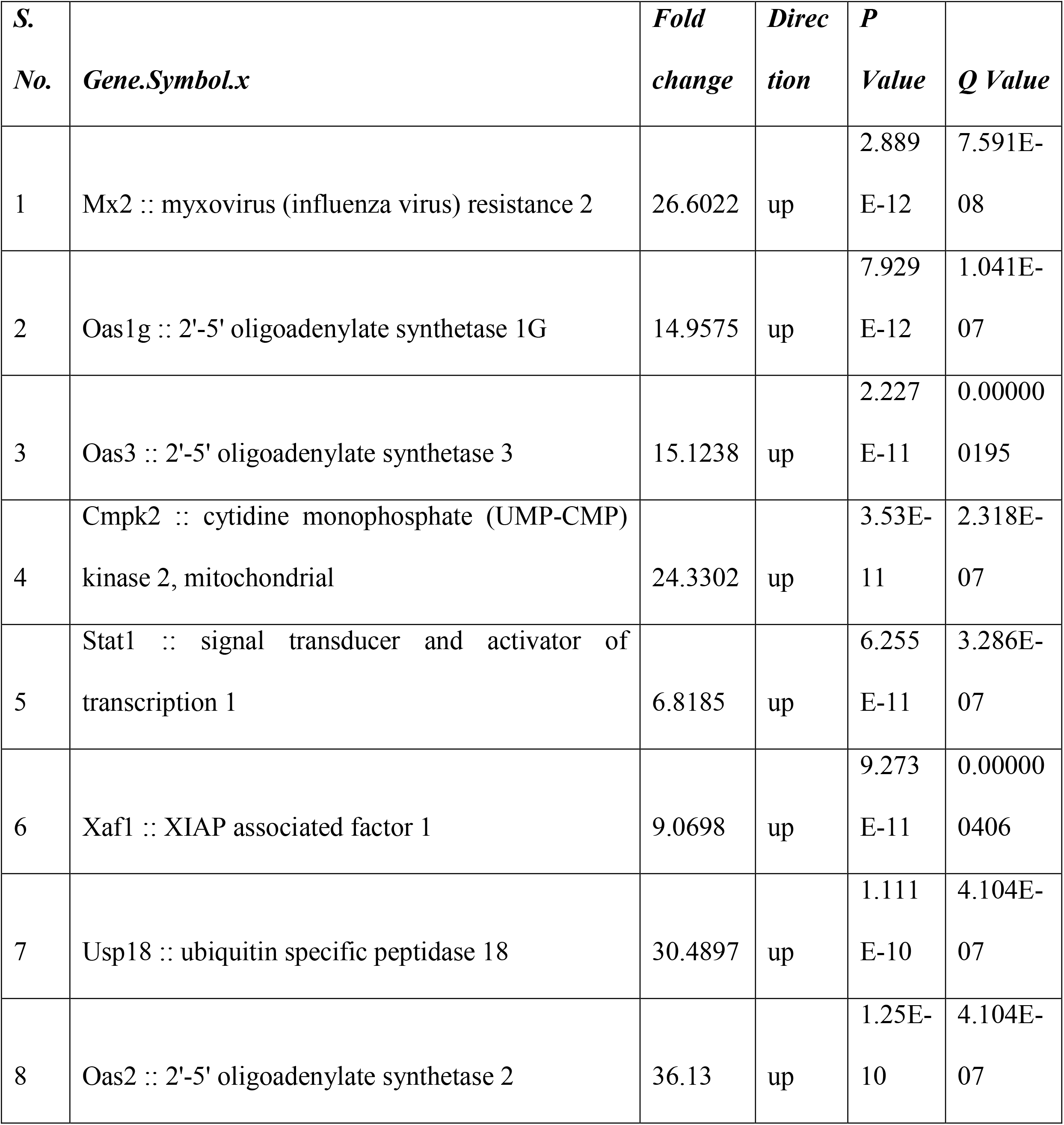
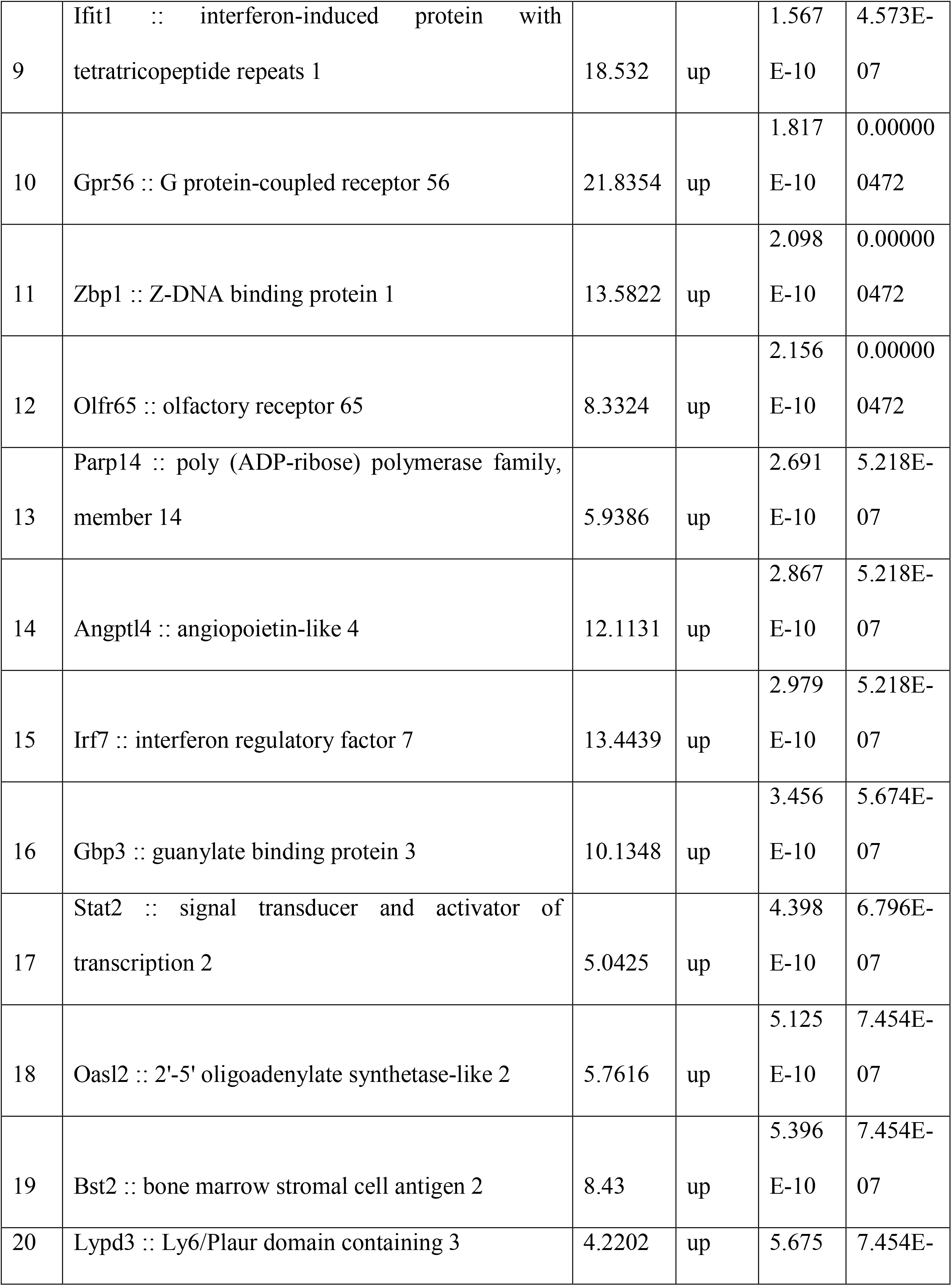
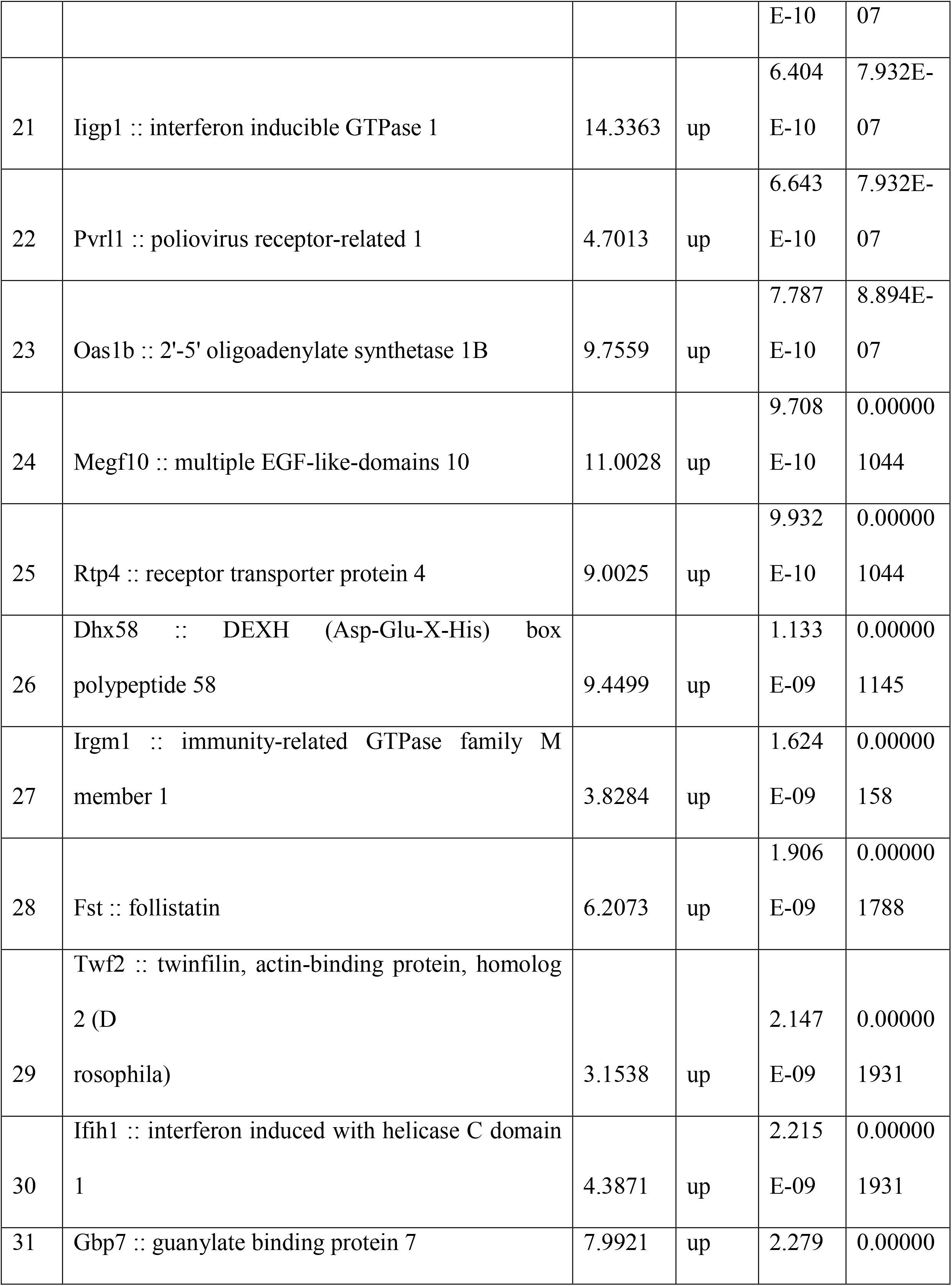
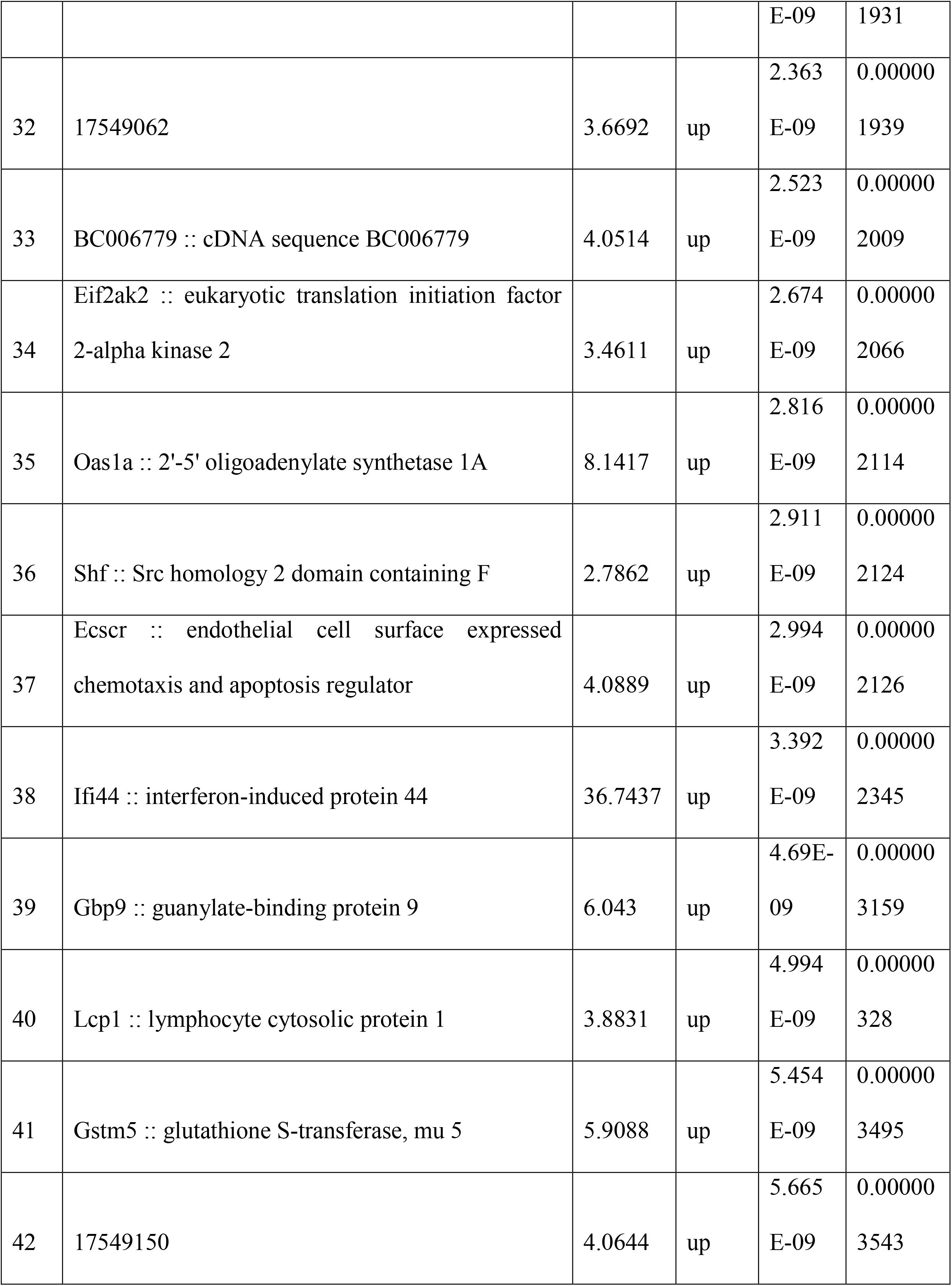
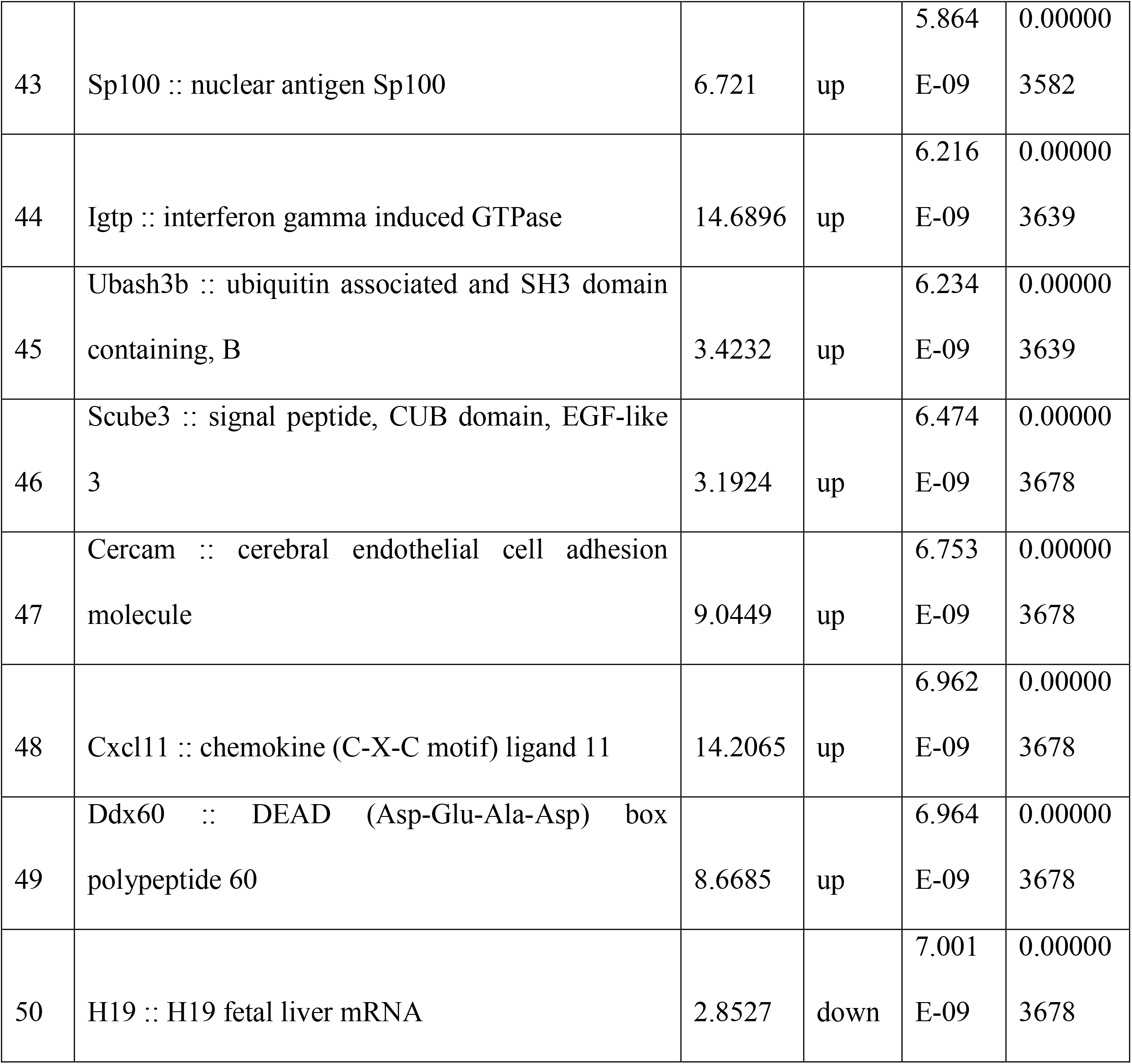
The top 50 DE genes generated from thegene expression profile of three independent replicates of control and Id KD cells was compared by microarray analysis to generate a list of differentially expressed genes between Id depleted and control cells.

**Supplementary Table 2.**
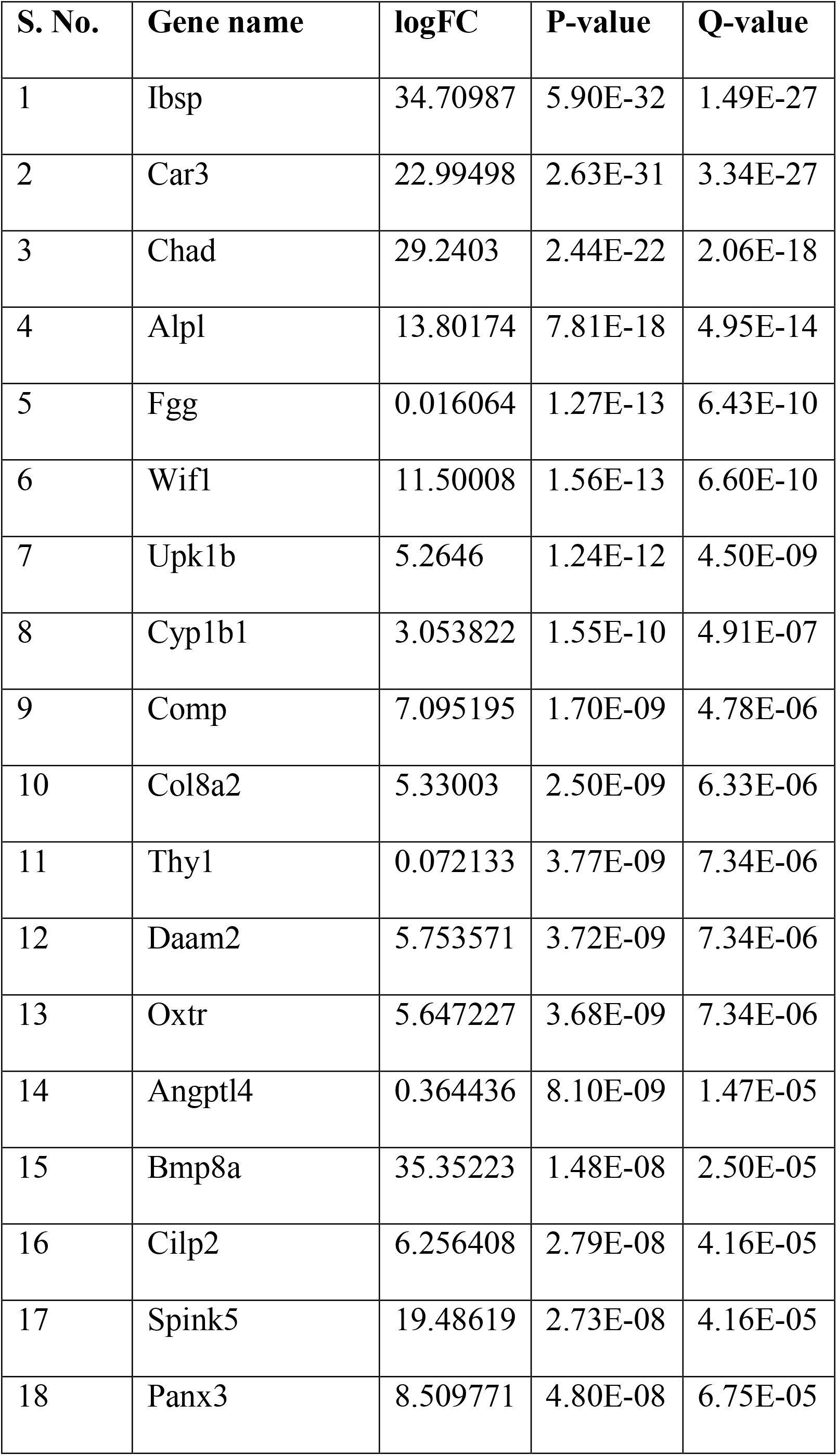
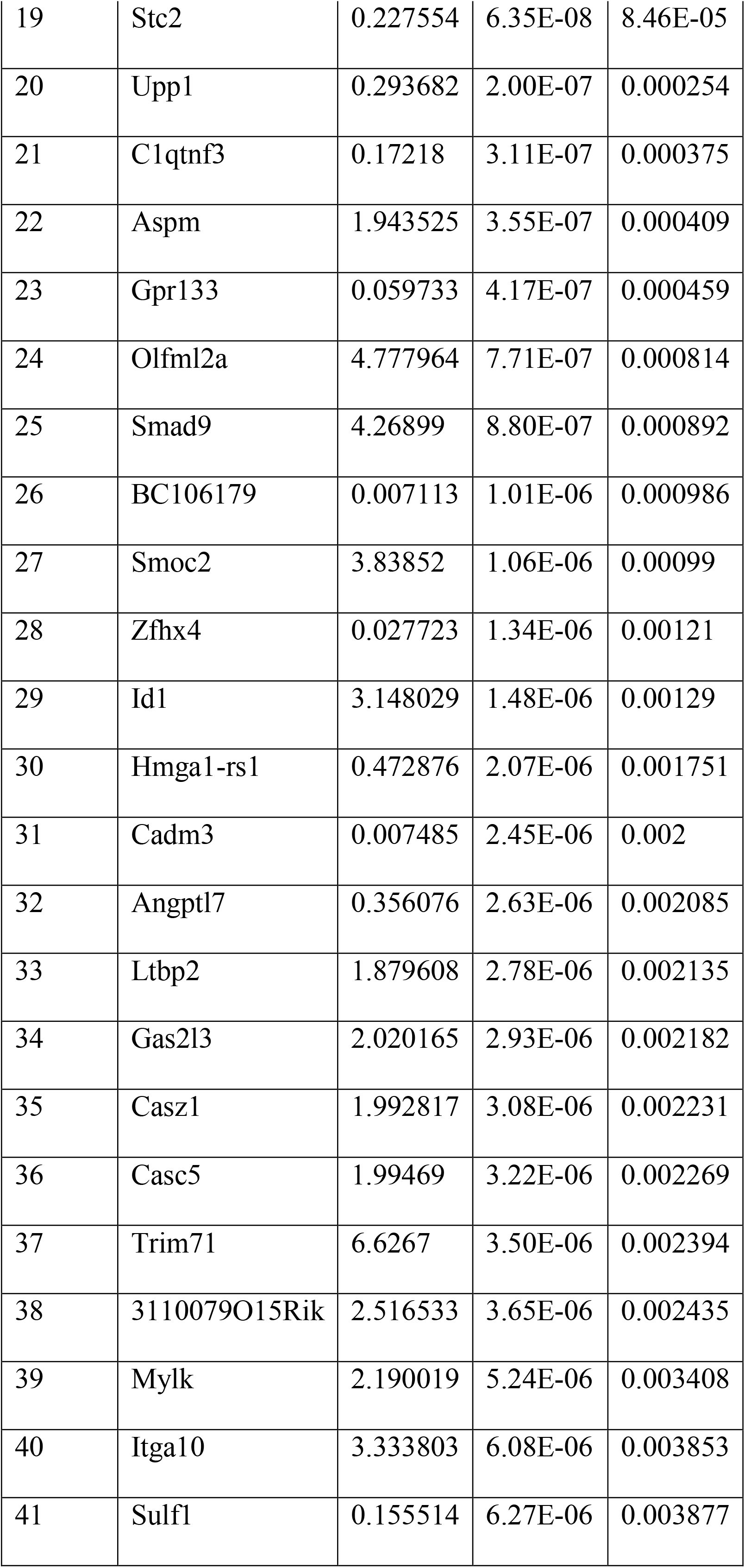
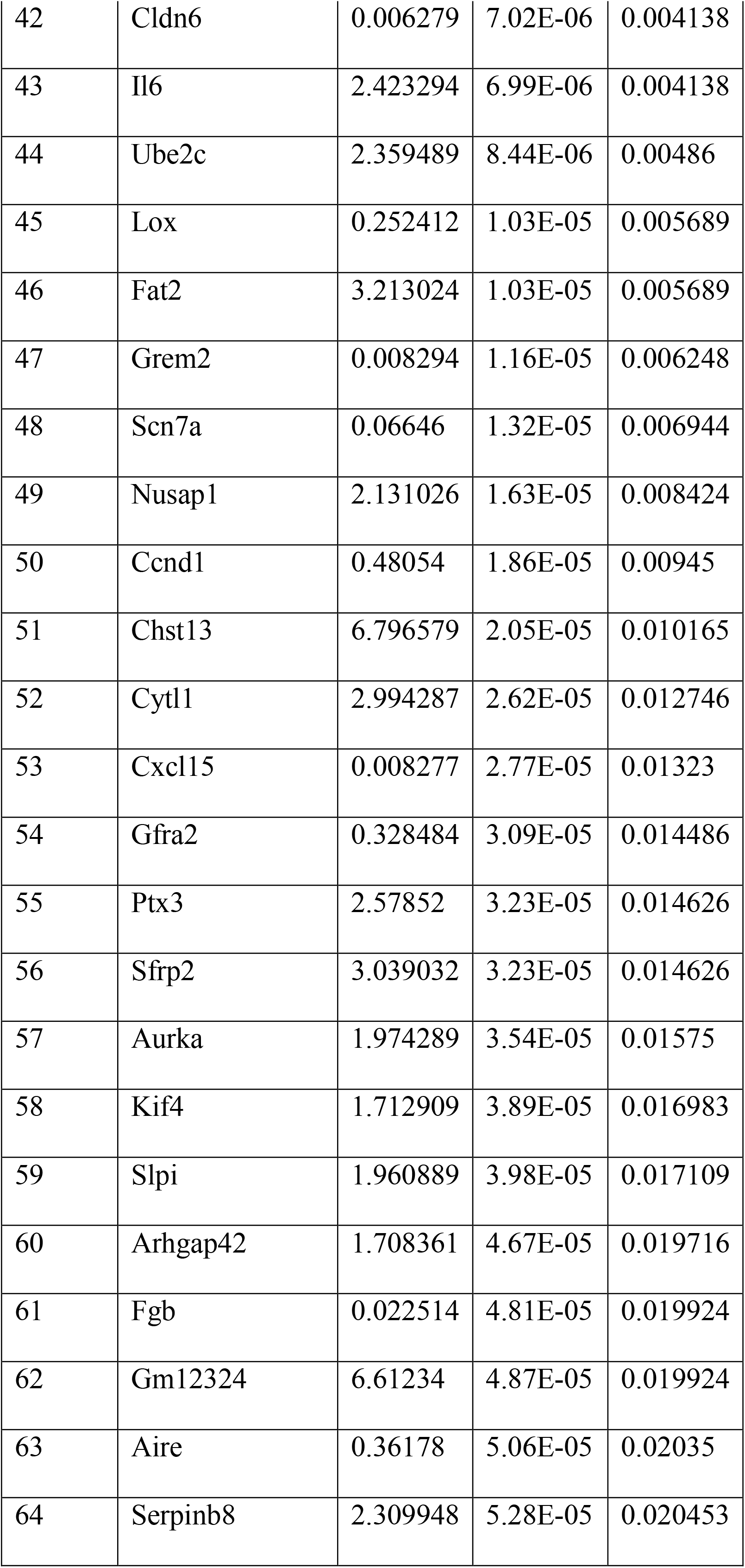
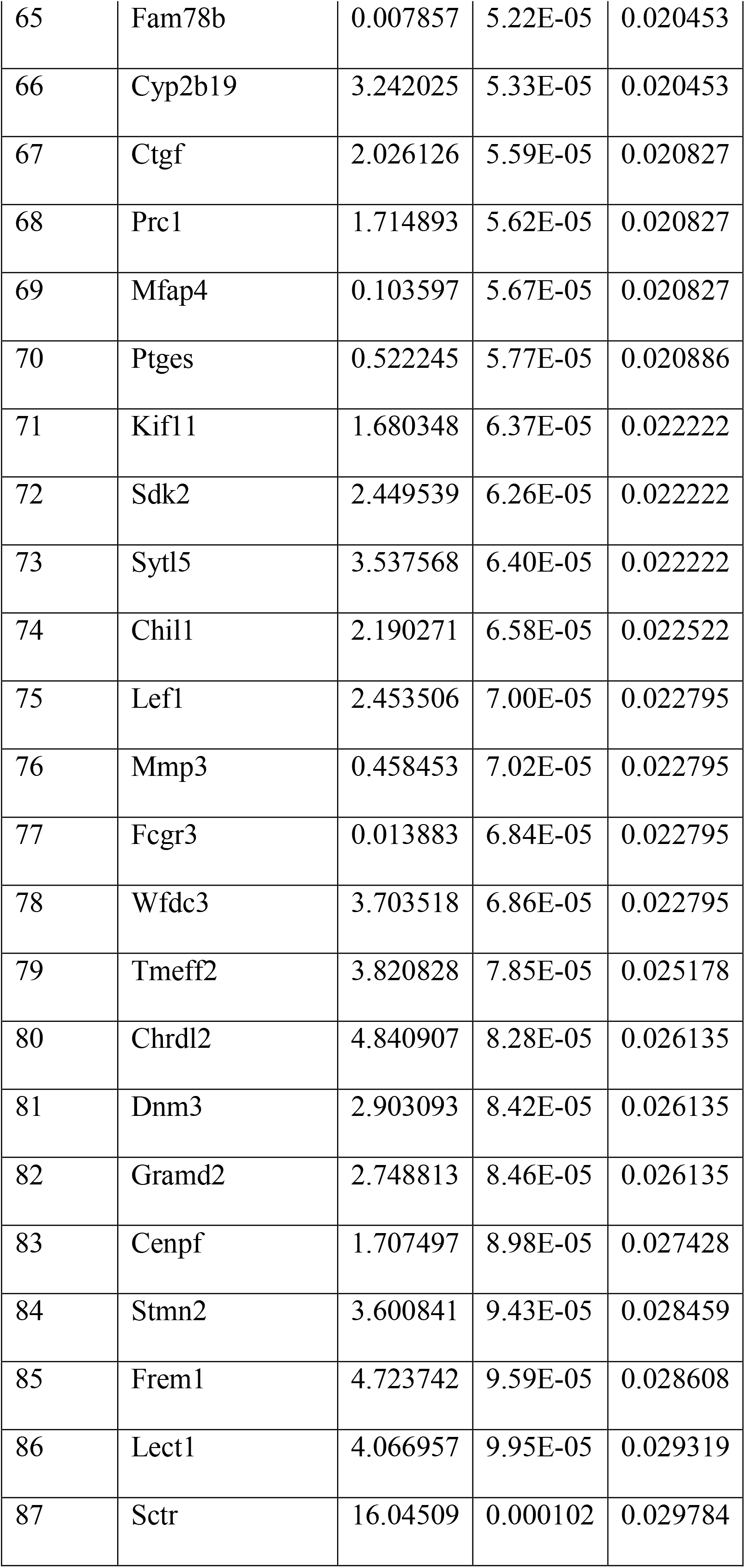
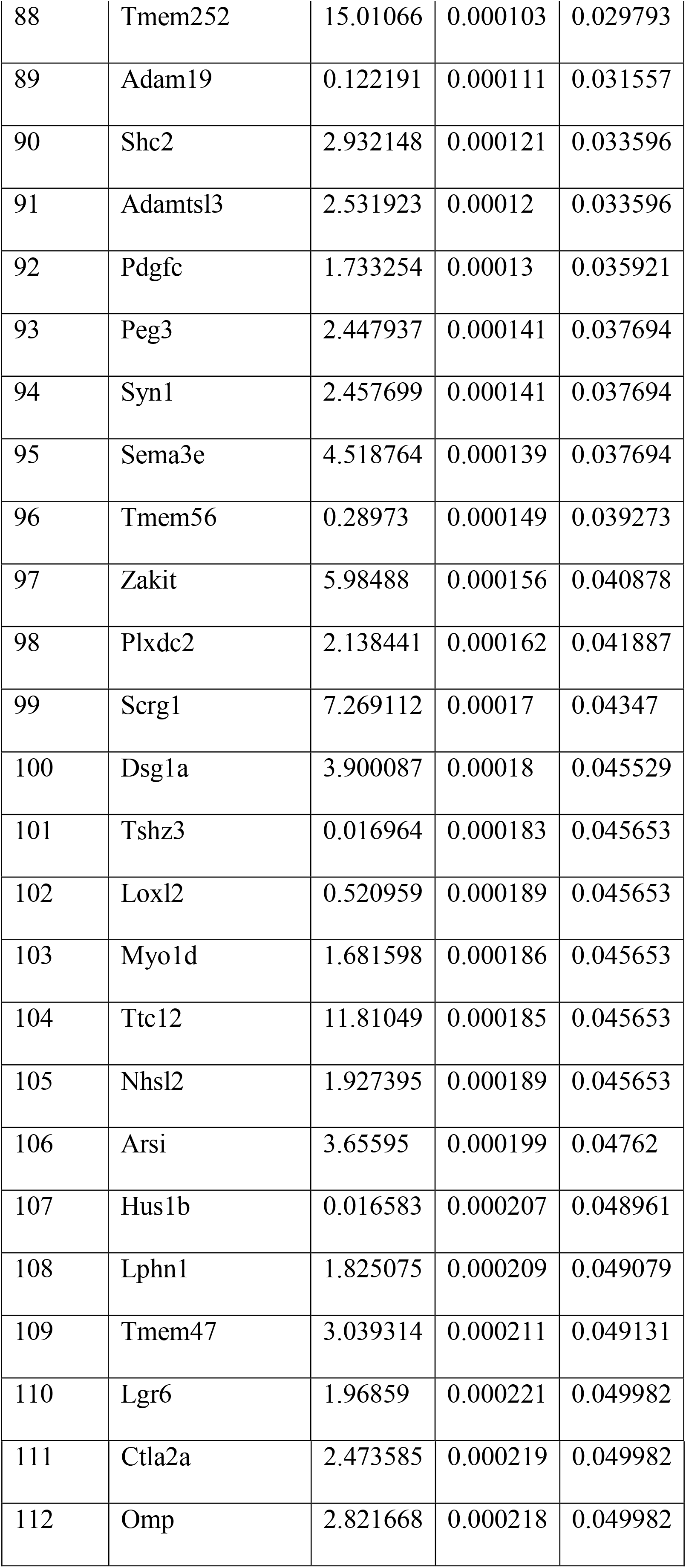
List of differentially expressed genes between the Id+ and Id-mouse TNBC cells generated from the Id1C3Tag model.

**Supplementary Table 3.** 61 candidate genes were identified for further validation as putative Id candidate target genes by siRNA screen.

| S. No. | miRNA.mimicsiRNA | Viability1 | Viability2 | Mean_Viability | Replicate_Disparity |
| --- | --- | --- | --- | --- | --- |
| 1 | Actb | 119.2385 | 69.44172 | 94.34012 | 0.199673 |
| 2 | Adamtsl3 | 59.5834 | 72.47348 | 66.02844 | 0.051686 |
| 3 | Alpl | 90.50029 | 96.10138 | 93.30084 | 0.022459 |
| 4 | ANGPTL4 | 47.11264 | 43.44723 | 45.27994 | 0.014697 |
| 5 | Aspm | 73.0994 | 49.45445 | 61.27693 | 0.094811 |
| 6 | AURKA | 47.13871 | 62.01435 | 54.57653 | 0.059648 |
| 7 | Axin2 | 120.1843 | 168.8596 | 144.522 | 0.195177 |
| 8 | BMI1 | 76.35547 | 93.35345 | 84.85446 | 0.068158 |
| 9 | BMP8A | 54.38554 | 78.88393 | 66.63473 | 0.098233 |
| 10 | Casc5 | 12.91458 | 20.8636 | 16.88909 | 0.031874 |
| 11 | CCND1 | 2.362772 | 2.487312 | 2.425042 | 0.000499 |
| 12 | Ccne1 | 107.1427 | 101.5924 | 104.3675 | 0.022255 |
| 13 | Cdkn1a | 151.815 | 187.5555 | 169.6852 | 0.143311 |
| 14 | Cdkn2a | 9.258506 | 12.23492 | 10.74671 | 0.011935 |
| 15 | CENPF | 104.4639 | 126.2672 | 115.3655 | 0.087426 |
| 16 | CHAD | 161.2562 | 192.6356 | 176.9459 | 0.125824 |
| 17 | CLDN6 | 5.437018 | 16.92079 | 11.1789 | 0.046047 |
| 18 | Col8a2 | 92.66478 | 91.96999 | 92.31739 | 0.002786 |
| 19 | Comp | 112.4995 | 166.9346 | 139.717 | 0.218272 |
| 20 | CXCL15 | 11.52211 | 22.00924 | 16.76568 | 0.042051 |
| 21 | Dpysl2 | 83.76739 | 73.22459 | 78.49599 | 0.042274 |
| 22 | FOXC2 | 32.42075 | 51.92788 | 42.17431 | 0.078219 |
| 23 | GAPDH | 74.66308 | 129.214 | 101.9385 | 0.218736 |
| 24 | Gp9 | 67.24151 | 72.35828 | 69.79989 | 0.020517 |
| 25 | GPR133 | 66.20689 | 72.52594 | 69.36641 | 0.025338 |
| 26 | GREM2 | 14.9732 | 27.7484 | 21.3608 | 0.051226 |
| 27 | Gypa | 55.29883 | 41.61613 | 48.45748 | 0.054864 |
| 28 | Ibsp | 56.76146 | 76.95146 | 66.85646 | 0.080957 |
| 29 | ID1 | 119.4619 | 135.7036 | 127.5827 | 0.065126 |
| 30 | ID2 | 25.04487 | 48.90737 | 36.97612 | 0.095683 |
| 31 | ID3 | 26.8322 | 26.7894 | 26.8108 | 0.000172 |
| 32 | Id4 | 60.58005 | 61.8837 | 61.23187 | 0.005227 |
| 33 | Il6 | 54.41804 | 38.86627 | 46.64216 | 0.062359 |
| 34 | Kif11 | 5.929065 | 10.06528 | 7.997174 | 0.016585 |
| 35 | Kif4 | 137.4777 | 145.0792 | 141.2785 | 0.03048 |
| 36 | Lef1 | 69.74422 | 75.35658 | 72.5504 | 0.022504 |
| 37 | Lgr6 | 13.77349 | 27.60068 | 20.68709 | 0.055444 |
| 38 | Lox | 30.49407 | 53.24569 | 41.86988 | 0.091229 |
| 39 | Loxl2 | 33.93647 | 41.46456 | 37.70052 | 0.030186 |
| 40 | Myom1 | 78.83341 | 66.06561 | 72.44951 | 0.051196 |
| 41 | Nusap1 | 26.12422 | 18.19966 | 22.16194 | 0.031776 |
| 42 | Oxtr | 14.45927 | 29.72044 | 22.08986 | 0.061194 |
| 43 | Panx3 | 70.37883 | 95.81948 | 83.09915 | 0.102011 |
| 44 | Pdgfc | 48.00148 | 46.44553 | 47.2235 | 0.006239 |
| 45 | POSTN | 57.7375 | 78.37966 | 68.05858 | 0.08277 |
| 46 | Prc1 | 44.28233 | 36.23709 | 40.25971 | 0.03226 |
| 47 | ROBO1 | 249.3912 | 225.3732 | 237.3822 | 0.096306 |
| 48 | Scrn1 | 219.7686 | 218.585 | 219.1768 | 0.004746 |
| 49 | Sctr | 102.5398 | 85.73719 | 94.13851 | 0.067375 |
| 50 | Sfrp2 | 113.2776 | 131.8863 | 122.582 | 0.074616 |
| 51 | Smad9 | 37.48249 | 43.25865 | 40.37057 | 0.023161 |
| 52 | Spaca3 | 46.28752 | 72.89763 | 59.59258 | 0.1067 |
| 53 | Stom | 174.221 | 165.451 | 169.836 | 0.035165 |
| 54 | TGFBR3 | 123.9611 | 98.65301 | 111.307 | 0.101479 |
| 55 | Tmem252 | 9.216349 | 8.13346 | 8.674905 | 0.004342 |
| 56 | Tmem47 | 33.99214 | 59.51196 | 46.75205 | 0.102328 |
| 57 | Ube2c | 66.20978 | 93.01105 | 79.61042 | 0.107467 |
| 58 | VCAM1 | 20.82949 | 33.1387 | 26.9841 | 0.049357 |
| 59 | Vps51 | 46.86967 | 34.84365 | 40.85666 | 0.048222 |
| 60 | WIF1 | 31.32273 | 52.26417 | 41.79345 | 0.08397 |
| 61 | ylk | 103.3257 | 62.85685 | 83.09127 | 0.162271 |

